# Succession-diagram-based Markov chains reveal the attractor landscape of asynchronous Boolean networks

**DOI:** 10.64898/2025.12.17.694936

**Authors:** Kyu Hyong Park, Réka Albert

**Affiliations:** Department of Physics, Pennsylvania State University, University Park, PA, USA

**Keywords:** Boolean network, Boolean model, Markov chain, attractor landscape, trap space, coarse-graining, stochastic dynamics, cell fate modeling

## Abstract

Comprehensive analysis of the dynamics of Boolean models of biological systems is hampered by the exponentially large state space. Here we introduce the succession-diagram-based Markov chain (SD Markov chain), a coarse-grained representation that uses trap spaces (unescapable state subspaces) of the Boolean model as the states of a Markov chain. These trap spaces and their succession diagram can be efficiently identified, and constitute a dramatic reduction compared to the full state space. The SD Markov chain preserves the decisions that trap the system’s dynamics while making the state space computationally tractable. Using an ensemble of random Boolean networks with known state transition matrices, we show that the SD Markov chain accurately reproduces attractors, basins of attraction, convergence probabilities, decision transitions, and sequences of events. By combining the interpretability of the succession diagram with the probabilistic rigor of Markov analysis, the SD Markov chain offers a compact quantitative description of the attractor landscape and provides a new avenue for studying control and stability in complex biological systems.

## Introduction

Biological systems consist of elements with distinct identities that interact in specific ways. In biomolecular systems, for example, each element may represent a gene transcript, protein, or small molecule, and their interactions include transcriptional regulation, protein-protein interactions, and chemical reactions. A well-established way to capture such element-specific interactions is the Boolean model^1–4^. Originally developed for electronic circuits^5^, Boolean modeling has been applied across biological scales, from gene regulatory networks^3,6,7^ and neuronal circuits^8^ to ecological and social communities^9,10^. In a Boolean model, each element is characterized by a binary state and a Boolean function that describes how the element responds to its regulators. The functions reflect the diversity of the elements and of their interactions. Boolean models are especially suited for biomolecular systems, which often feature nonlinear, sigmoidal regulation^11,12^. Boolean models are the simplest among dynamic models: they have no parameters and characterize each element with the fewest states.

The model’s global state is the vector of all element states, and its dynamics unfold through discrete state transitions. In the frequently-used stochastic asynchronous update scheme, a randomly selected element is updated at each time. This update scheme aims to reflect variations in or lack of knowledge regarding the durations of the processes described by the model^13^. Boolean models of specific biological systems successfully reproduced key system outcomes. For example, Boolean models of biomolecular networks reproduced cell phenotypes and behaviors^7,14–17^. Boolean models also yielded many useful predictions, such as identifying master regulators or drug targets^14,18,19^. Predictions derived from Boolean models of various biological systems were verified experimentally^19–22^.

A key strength of dynamic models lies in predicting the modeled system’s behavior in unexplored conditions. Because Boolean models have a finite number of states, it is in principle possible to map their whole state space and find out every long-term behavior of the system. The state transition matrix provides a comprehensive summary of the probability of each elementary transition. This information provides the building blocks of every system behavior (i.e., every trajectory in state space). In a Boolean model, each state change depends on the starting state only, thus satisfying the Markov property^23^. Thus, in principle Markov chain theory can be used to determine the probability of each long-term behavior. However, the practical challenge is that the number of system states increases exponentially with the number of elements. For a system with 100 elements the state transition matrix contains 2^200^ elements, far beyond the current computational and storage capacity.

To bypass this challenge, modelers often sample trajectories by employing simulations, e.g., utilizing one of multiple software tools^24–27^. Simulations have proven useful for determining average behaviors^28^. Yet, they can only cover a tiny fraction of the state space, making it likely that dynamical behaviors are missed. An effective alternative method to determine the repertoire of long-term behaviors of a Boolean model is via the identification of so-called stable motifs^29^. A stable motif is a self-sufficient feedback loop that can sustain an associated state of its constituent elements regardless of the rest of the system. The locking-in of a stable motif reduces the freedom of the system, enclosing it into a subset of the state space called a trap space^30^. Trap spaces nest within each other, and inclusion-minimal trap spaces correspond to attractors^31^. Locking in one stable motif versus another is an irreversible decision that selects a trap space and channels the system toward an attractor.

The inclusion relations of trap spaces form a directed acyclic graph, called a succession diagram, which provides a complete summary of all the decisions possible in the model. The number of trap spaces in the succession diagram is much lower than the number of states. For example, the ensemble of 212 Boolean models of biological systems collected in the Biodivine Boolean Model (BBM) repository^32^ has a median of 38 elements. A 38-variable Boolean model has around 300 billion states. In contrast, analysis of the succession diagrams of this model ensemble^31^ yielded a median succession diagram size of 87, which is more than nine orders of magnitude lower than the size of the state space. Succession diagrams are both tractable and interpretable, offering a qualitative view reminiscent of Waddington’s epigenetic landscape^33^. Yet, a limitation of the succession diagram is that it does not describe the trajectories of the system, nor the probability of selecting a decision among multiple possibilities.

Here we introduce the succession-diagram-based Markov chain (SD Markov chain), which combines the interpretability of the succession diagram with the quantitative rigor of Markov chain theory. The states of this Markov chain are the trap spaces of the Boolean model. Thus, the state space of the SD Markov chain is tractable, making it possible to construct the state transition matrix. To quantify the loss of information arising from state grouping, we perform a thorough evaluation of the SD Markov chain for an ensemble of Boolean models for which the state transition matrix can be identified. We find that the SD Markov chain is remarkably accurate in capturing the attractors, their reachability from arbitrary initial conditions, the decision transitions, and the sequences of events.

## Background and key concepts

### Foundations of Boolean modeling

Boolean dynamical systems are often referred to as Boolean networks. A Boolean network consists of an interaction graph, the Boolean functions of each element, and the update scheme. When a Boolean network is used to describe a real-world complex system, we will refer to it as a Boolean model.

The nodes of the interaction graph represent the elements of the system, and its directed edges represent regulatory influences. Signs are often assigned to the individual edges, indicating an activating (+1) or inhibiting (−1) influence. Each node of a Boolean network can take on one of two states at any given time step, ON (1) or OFF (0). The node state is updated according to a Boolean function of the states of the node’s regulators. The update scheme specifies how nodes are selected for update in each time step^28^. The node update induces an update of the system state, called a state transition. Here we use stochastic asynchronous update, which is a versatile choice that avoids certain shortcomings associated with synchronous update^2,28^. The standard practice is to assume that each node has the same probability of being selected for update.

Each Boolean network with *N* nodes induces a state transition graph (STG) whose 2^*N*^ nodes represent all possible system states. A directed edge from a system state *s*_*i*_ to another state *s*_*j*_ exists if and only if *s*_*i*_ can be updated in one time step to obtain *s*_*j*_. A more complete information is offered by the state transition matrix, of size 2^*N*^ × 2^*N*^, which for any pair of states *s*_*i*_, *s*_*j*_ indicates the probability that updating state *s*_*i*_ leads to state *s*_*j*_ ^34^. Under stochastic asynchronous update, each state has a maximum of *N* transitions, thus the maximal density of edges in the state transition graph is *N*/2^*N*^, which makes the STG very sparse. This property motivates coarse-graining of the STG into a hierarchical transition graph^35^, which can be constructed more effectively (but still with exponential complexity) than the full STG^36^.

The attractors of a Boolean network are the smallest sets of states that are invariant under the network’s dynamics (i.e., in which the system is trapped)^1,2^. Each point attractor consists of a single state, and can be identified as a terminal node of the STG. Each complex attractor contains multiple states that the system keeps revisiting, and can be identified as a terminal strongly connected component of the STG. States that do not belong in an attractor are referred to as transient states. The states that belong to or can reach an attractor via state transitions make up the basin of attraction of the attractor. States that belong in the basin of a single attractor are said to belong in the strong basin of that attractor. States that reach multiple attractors belong to the weak basin of those attractors.

### Stable motifs, trap spaces, and the succession diagram

A stable motif of a Boolean network consists of a positive circuit of the interaction graph and an associated state of the constitutive nodes. The key property of the stable motif is that once its nodes attain this state, the inter-regulation of the nodes is able to maintain this state, regardless of the dynamics of the rest of the network. Once locked in, a stable motif is irreversible and traps the system into a trap space, i.e., a subspace with a set of fixed node values that cannot be escaped^30,37^.

The successive locking-in of stable motifs traps the system into smaller and smaller trap spaces. Iterative identification of stable motifs followed by network reduction (percolation of constants) results in a graph called a succession diagram^29,37^. The succession diagram is a directed acyclic graph whose nodes correspond to trap spaces and whose edges indicate inclusion. An important feature of the succession diagram is that any trajectory of the Boolean network that does not follow a path of this directed acyclic graph is forbidden. For example, no trajectory can exist that goes from a trap space *T*1 to a less restrictive trap space *T*2 ⊃ *T*1, or to a trap space *T*3 that does not intersect *T*1.

The leaf nodes of the succession diagram are minimal trap spaces. A minimal trap space is either a single state (a point attractor) or a set of states in which the nodes that are not fixed by stable motifs have unspecified states and may oscillate between the 0 and 1 state. A minimal trap space is a good approximation of an attractor. A minimal trap space rarely contains more than one attractor, and attractors that lie outside of minimal trap spaces are extremely rare^38,39^.

### Foundations of Markov chains

A Markov chain is a stochastic process wherein the probability of reaching a future state only depends on the current state^23,40,41^. The Markov chains of interest here are discrete-time Markov chains with a countable number of states. Such a Markov chain is defined as a set of states and a matrix describing the transition probabilities between these states. The transition matrix is stochastic, meaning that each row represents probabilities that sum up to 1. Markov chain theory can be used to determine which states are transient and which are recurrent. The recurrent states form one or more recurrent classes. The states of a recurrent class only have transitions to each other, and do not have transitions that leave the class. Thus, each recurrent class forms a block in the transition matrix raised to an infinite power. Absorbing states are a special type of recurrent class where this block is of size 1. Absorbing states can also be identified in the transition matrix itself as states for which the probability of staying the same is 1 and all other transition probabilities are 0. Markov chain theory indicates how to calculate the probability of convergence into a specific state (e.g., an absorbing state) from an arbitrary initial state and the expected number of steps prior to convergence^23,40,41^. Markov chains are used in diverse fields, including as models of the conformational dynamics of biomolecules^42^ and the population dynamics of cells^43^.

### Characterizing stochastic asynchronous Boolean networks with Markov chains

In a stochastic asynchronous Boolean network, the next state of the system after the update of a node depends only on the current state of the node’s regulators. This property makes the Boolean network a Markov chain^34^. For a Boolean network of *N* nodes, the states of this Markov chain are the 2^*N*^ states of the Boolean network. The transition matrix contains a nonzero probability for every state transition allowed by the update function of a node. In the standard implementation of stochastic asynchronous update, the transition probability is 1/*N* for each allowed transition between different states. The probability of staying in the same state is 1/*N* multiplied by the number of nodes whose update leaves the state unchanged.

The recurrent classes of the Markov chain correspond to the attractors of the Boolean network. Absorbing states are point attractors. Each class of multiple recurrent states corresponds to a terminal SCC in the STG, and thus to a complex attractor. The various powers of the transition matrix indicate the transition probability of any state into any other state in as many time steps. In particular, the Markov chain indicates the probability of convergence into an absorbing state (point attractor) or a recurrent state that is part of a complex attractor. These probabilities offer a quantitative description of the attractor landscape. Unfortunately, the exponential scaling of the transition matrix’s size ( 2^2*N*^) makes Markov chain analysis difficult for systems with many nodes.

### Defining the transition probabilities between groups of states

The key idea pursued here is to describe a stochastic asynchronous Boolean network by a coarse-grained Markov chain whose states correspond to groups of states in the Boolean network. In this work, we consider that the state transition graph (STG) of the Boolean network is known. We use this STG both to construct the transition probabilities in the Markov chain, and to evaluate the predictions that arise from the Markov chain. In this subsection, we describe the methodology of obtaining the transition probabilities of the coarse-grained Markov chain from the transition probabilities of the Boolean network, and the methodology of translating the transition probabilities of the Markov chain into approximate transition probabilities in the Boolean network. We illustrate these methodologies on the Boolean network of Example **1**, which we will use throughout this work.

Consider that we are given a mapping π: *S* → *C* between the state space *S* of the Boolean network and the state space *C* of the coarse-grained Markov chain such that π(*s*) = *c* for *s* ∈ *S* and *c* ∈ *C*. A state *c* of the Markov chain corresponds to a group of states *G* = π^−1^ (*c*) = {*s* ∈ *S*|π(*s*) = *c*} of the Boolean network. For simplicity, we use the same index when a Boolean state corresponds to a Markov state, i.e., *s*_*i*_ ∈ *G*_*i*_ = π^−1^ (*c*_*i*_).

We illustrate the state assignment of a coarse-grained Markov chain using the Boolean network of Example **1**. The 8 states of the Boolean network are {000, 001,.. 111}, where each system state indicates the state of nodes **A, B**, and **C** in alphabetical order. The coarse-grained Markov chain has six states, *c*_1_ = π(101) = π(110), *c*_2_ = π(010) = π(001), *c*_3_ = π(100), *c*_4_ = π(111), *c*_5_ = π(000) and *c*_6_ = π(011). For ease of interpretation of all the analyses of this system, we avoid using two sets of notations and instead refer to the states of the coarse-grained Markov chain by the corresponding groups of Boolean states. Namely, we introduce the notations *G*1 = π^−1^(*c*_1_) = {101, 110} and, *G*2 = π^−1^ (*c*_2_) = {010, 001} and instead of *c*_1_, *c*_2_, *c*_3_, *c*_4_, *c*_5_, *c*_6_, we use *G*1, *G*2, 100, 111, 000, 011, respectively.

#### Example 1

In a three-node Boolean network, **B** and **C** form a positive cycle, and **A** is regulated by **C** and itself. The update functions are

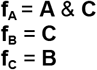

Given π, the transition probabilities in the Boolean network can be used to define the transition probabilities of the Markov chain. Specifically, the transition probability between a Markov state *c*_*j*_ and a Markov state *c*_*k*_ is the sum of the probabilities of all the transitions *s*_*j*_ → *s*_*k*_ (i.e., all the transitions from *G*_*j*_ to *G*_*k*_) in the Boolean network, divided by |*G*_*j*_| to make the transition probabilities in a row of the Markov chain add up to 1.

We illustrate this process in Figure **1**, which uses the Boolean network of Example **1** and the coarse-grained Markov chain defined earlier. The first matrix is the transition matrix of the Boolean network. The submatrix that corresponds to transitions from Boolean states in *G*1 to Boolean states in *G*2 is highlighted in yellow. The second matrix is the transition matrix of the coarse-grained Markov chain. The transition probability from *G*1 to *G*2, *p*_*G*1*G*2_, is ⅓ (the only nonzero entry in the yellow-highlighted submatrix) divided by |*G*1| = 2.

**Figure 1.**
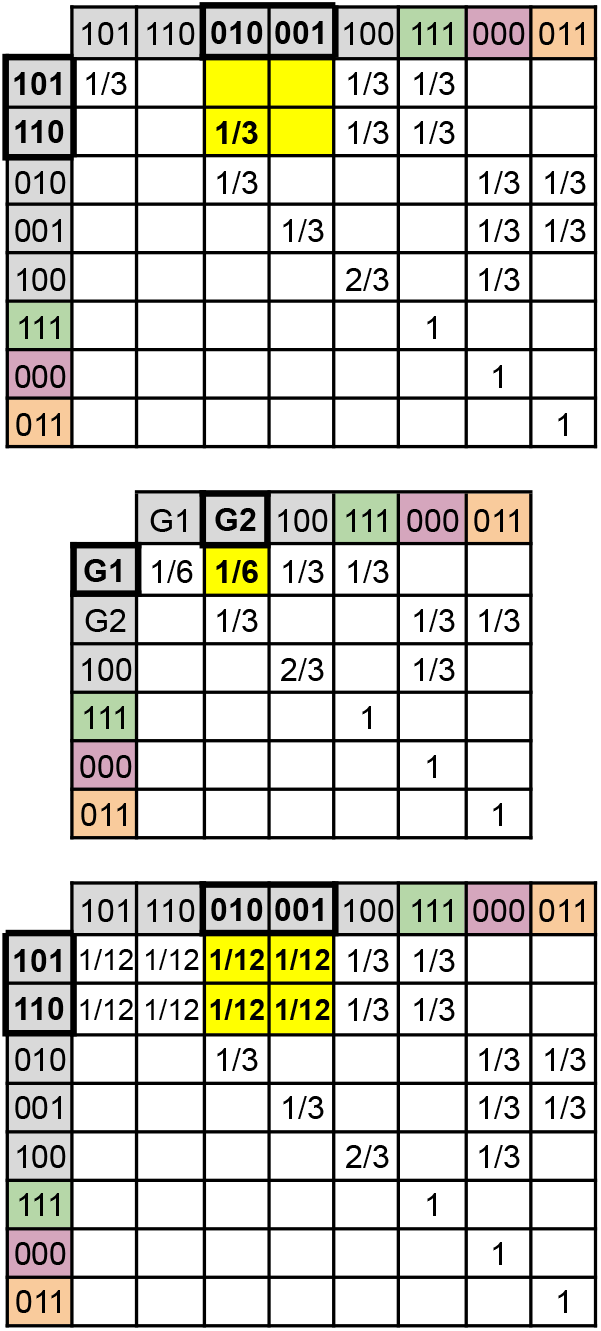
The transition matrix of a Boolean network (top), the transition matrix of a Markov chain of grouped states (middle), and the re-expanded transition matrix that serves as the approximation of the original (bottom). The states are indicated on the left and at the top. Attractor states are color-coded. The groups of states G1={101, 110} and G2={010, 001} form corresponding states of the Markov chain. The transition probability of the Markov chain from G1 to G2 (highlighted in yellow) is obtained by taking the sum of the 2 x 2 submatrix of the Boolean network representing all transitions among G1 and G2 (also highlighted in yellow) and dividing by 2 (the size of G1). The Markov chain’s transition matrix can be re-expanded to approximate the state transition probabilities of the Boolean network. The transition from G1 to G2 will be expanded to a 2 x 2 block in the re-expanded transition matrix (highlighted in yellow). The expansion assumes uniform probability for the transitions that lead to the same group.

The more states become grouped, the smaller the coarse-grained Markov chain becomes, and the more feasible it is to analyze it. After processing the smaller transition matrix with certain methods, such as raising it to the infinite power, the transition probabilities can be translated back to the Boolean network. During this process, each coarse-grained transition is evenly distributed among its constituent states. Thus, a transition probability *p*_*jk*_ between a Markov state *c*_*j*_ and a Markov state *c*_*k*_ will be expanded to |*G*_*j*_ |× |*G*_*k*_ |entries of the re-expanded transition matrix, each having a probability of *p*_*jk*_ /|*G*_*k*_ |. Importantly, the differences between the members of the same group disappear. To illustrate this fact, in Figure **1** we present the coarse-grained and then re-expanded transition matrix of Example **1**. The transition probability between Markov states *G*1 and *G*2 is equally distributed to the four corresponding pairs of Boolean states. This equal distribution differs from the Boolean network’s transition matrix, in which only one transition had a nonzero probability. It is generally true that the re-expanded transition matrix approximates the original transition matrix with a block structure that predicts more possible transitions than the original and never fewer.

### Defining succession-diagram-based Markov chains

The key innovation of our approach is to define the states of the SD Markov chain as the trap spaces of the succession diagram of the Boolean network. Such a Markov chain has a state space that is many orders of magnitude smaller than the state space of the Boolean network. To estimate the state-space reduction, we followed up a succession diagram analysis of the BBM model ensemble with fixed inputs^31^ (see the Methods for details). The ensemble has 129 models whose state space is larger than 1 million states. Yet, 109 of these models have 20 or fewer trap spaces in the succession diagram. These models are not amenable to STG analysis but are eminently tractable by SD Markov chain approaches.

Traditional Markov chain lumping approaches^44,45^ require exact equivalence of transition probabilities for every state that is assigned to the same group. The states in the same trap space do not usually have the same transition probabilities, but they have a very important commonality: they agree that transitions that would escape the trap space are impossible. In the following, we define SD Markov chains and illustrate them in an example.

Each state of a *succession-diagram-based Markov chain* corresponds to the states in a trap space of the Boolean network that are not contained in a smaller trap space. Figure **1** illustrates the SD Markov chain of Example **1**. As an example, there are four states in the trap space 0**, namely {000, 001, 010, 011}. Since 011 and 000 also belong to smaller trap spaces (which are the states themselves), only the two states {001, 010} form the group G2 that corresponds to the trap space 0**. Note that each minimal trap space of the Boolean network becomes a separate absorbing state of the SD Markov chain. The nested trap space structure ensures that the grouped states agree on forbidden transitions. For example, the succession diagram of Example **1** shows that any transition from states in 0** to state 111 is impossible. By grouping states within 0** (G2 in this case), the grouped states all agree on not having a transition to 111. If we were to group states from two different trap spaces (e.g. from *11 and 0**), this property would not hold.

It is important to note that the hierarchical nature of the succession diagram makes the SD Markov chain a directed acyclic graph with self-edges. In addition to the self-edges of the absorbing states, self-edges arise from transitions among Boolean states mapped to the same state of the SD Markov chain. Indeed, all the states of the SD Markov chain of Example **1** have self-edges. The lack of cycles means that the only type of recurrent states in the SD Markov chain are absorbing states. A practical benefit of the acyclic nature of the SD Markov chain is that it allows the enumeration of all possible paths.

#### Alternative coarse-grained Markov chains

When evaluating the fidelity of the SD Markov chain in reflecting the properties of the Boolean system, it is useful to consider alternative coarse-grained Markov chains as benchmarks.

Our first benchmark Markov chain shares the SD Markov chain’s good feature of making each minimal trap space of the Boolean network into a state of the Markov chain. This Markov chain merges all the Boolean states outside of minimal trap spaces (states that are therefore transient) into a single state. We call this Markov chain the *Null Markov chain*. The Null Markov chain has fewer states than the SD Markov chain, thus it is more feasible to analyze. Its downside is that its merging of all transient Boolean states into a single Markov state makes it unable to distinguish basins of attraction or capture transient behaviors of the Boolean network.

To consider an alternative grouping that yields the same number of states as the SD Markov chain, we also define an ensemble of *Random Markov chains*. Each such Random Markov chain contains each minimal trap space of the Boolean network as a separate state. The rest of the states of the Boolean network are distributed uniformly randomly into groups in such a way that the number of states in the Random and SD Markov chains is the same. Because each transient state of a Random Markov chain contains a randomly selected mix of transient Boolean states, the Random Markov chain contains more transitions between pairs of transient Markov states, and from transient states to absorbing states, than the SD Markov chain. In contrast to the acyclic SD Markov chain, the transient states of the Random Markov chains form almost complete subgraphs, as we describe in the next section. We generate 100 Random Markov chains and take an ensemble average when comparing them with other methods. Figure **1** illustrates the transition network of the SD Markov chain, the Null Markov chain, as well as one example of a Random Markov chain. While the absorbing states are shared by all three types of Markov chains, they differ in their ability to capture attractor reachability from transient states, as we will show in later sections.

### Summary of the evaluation of the SD Markov chain

The SD Markov chain can predict multiple aspects of the dynamics of the Boolean network in an explanatory way. We perform a series of analyses on a spectrum from qualitative categorizations, whose results are easy to interpret and compare between systems, to quantitative probabilities, whose interpretation is more complex. Most of the analyses focus on characterizing the attractor landscape, i.e., the identity of the attractors and the reachability of attractors from arbitrary initial states. The final analysis evaluates the correspondence of paths in the SD Markov chain with trajectories of the Boolean network.

We use an ensemble of random Boolean networks (RBNs) with 10 nodes as a benchmark ensemble. This small network size allows the determination of the state transition graph and transition probabilities. The details of these Boolean networks are described in the Methods. The SD Markov chains constructed from this random ensemble have between 1 and 68 states, 7.6 being the average. This is a significant reduction from the 1024 states of 10-node Boolean networks. For each system, the Boolean state transition probabilities serve as the ground truth. We determine the succession diagram of the Boolean network and determine the states and transition probabilities of the SD Markov chain, Null Markov chain, and Random Markov chains.

To ground the analyses that follow, we first look into the network structure of the three Markov chains (see Methods for details). Of particular interest is the difference in the connectivity of the SD Markov chain and the Random Markov chain, which have the same number of states by design. We find that the Random Markov chains are close to having the maximal number of transitions that a Markov chain with a given number of transient states and absorbing states can have (see Methods for the formula), whereas the SD Markov chain is much more sparse. To quantify the difference, we compare the ratio of the number of transitions in each Markov chain and the number of transitions in the corresponding maximally-connected Markov chain. Intuitively, such a maximally-connected Markov chain is made up of a complete subgraph of transient states and has a transition between every transient state and every absorbing state. The median of the SD Markov chains’ ratio is 0.52. The average ratio of the 100 Random Markov chains has a median of 0.94. In conclusion, the Random Markov chains have nearly maximal connectivity. The acyclic structure of the SD Markov chain leads to a significantly sparser network of transitions, which is a closer reflection of the Boolean system’s sparse STG.

We calculate multiple properties of the attractor landscape of each type of Markov chain and compare them to the corresponding property of the Boolean network. The comparison involves metrics based on qualitative comparisons, namely precision, recall, negative predictive value (NPV), and specificity, and the quantitative metrics of root mean square difference (RMSD) and Kullback-Leibler Divergence (KLD), as described in the Methods. The ideal value for the qualitative metrics is 1, and the ideal value of RMSD and KLD is 0. Table **1** summarizes the key results of these analyses. Across all evaluated metrics, the SD Markov chain reproduces the attractor landscape of the Boolean networks with high precision and minimal information loss. We present each analysis in detail in the sections that follow.

**Table 1.** Summary of qualitative and quantitative metrics describing the predictive power of the SD Markov chain. The analyses are done on the full RBN ensemble or the most appropriate subset, and the values indicate the averages over the ensemble. The results of each SD Markov chain are compared with the corresponding results for 100 Random Markov chains and the Null Markov chain. We also indicate the results for suitable references described in the last column. The gray font indicates results that arise from the design of a coarse-grained Markov chain or of the references. The paths of Random and Null Markov chains are not comparable to the trajectories of the Boolean system.

| Metrics | SD MC | Random MC | Null MC | Reference | Reference description |
| --- | --- | --- | --- | --- | --- |
| Attractor vs non-attractor states | Prec.=0.97<br>Recall=1<br>NPV=1<br>Spec.=0.93 | Same as SD MC | Same as SD MC | Prec.=0.21<br>Recall=1,<br>NPV undefined<br>Spec.=0 | All states are attractor states |
| Strong and weak basins | Prec.=1<br>Recall=0.86<br>NPV=0.93<br>Spec.=1 | Prec.=1<br>Recall=0.30<br>NPV=0.58<br>Spec.=1 | Prec.=1<br>Recall=0.30<br>NPV=0.58<br>Spec.=1 | Prec. undefined<br>Recall=0<br>NPV=0.48<br>Spec.=1 | All states are in weak basins |
| Attractor reachability | Prec.=0.97<br>Recall=1<br>NPV=1<br>Spec.=0.90 | Prec.=0.74<br>Recall=1<br>NPV=1<br>Spec.=0.28 | Prec.=0.74<br>Recall=1<br>NPV=1<br>Spec.=0.27 | Prec.=0.64<br>Recall=1<br>NPV undefined<br>Spec.=0 | All attractors are reachable from any state |
| Convergence probabilities | RMSD=0.01<br>KLD=0.12 | RMSD=0.26<br>KLD=0.50 | RMSD=0.26<br>KLD=0.51 | RMSD=0.35<br>KLD=0.64 | Equal convergence to each attractor |
| Basin fractions | RMSD=0.05<br>KLD=0.04 | RMSD=0.13<br>KLD=0.13 | RMSD=0.13<br>KLD=0.13 | RMSD=0.12<br>KLD=0.09 | Equal convergence to each attractor |
| Average node values | RMSD=0.05 | RMSD=0.10 | RMSD=0.10 | RMSD=0.30 | Node average=0.5 |
| Decision transitions | Prec.=0.91<br>Recall=0.81<br>NPV=0.98<br>Spec.=0.99 | Prec.=0.41<br>Recall=0.17<br>NPV=0.91<br>Spec.=0.97 | Prec.=0.41<br>Recall=0.17<br>NPV=0.91<br>Spec.=0.96 | – | – |
| Fidelity of paths | Recall=1<br>Prec.=1.00 | Not informative | No paths | – | – |
| Path probabilities | RMSD=0.08 | Not informative | No paths | RMSD=0.16 | The nonzero transition probabilities are randomized |

### The SD Markov chain captures attractors

A recurrent class of a coarse-grained Markov chain corresponds to an attractor of the Boolean network. This is because by its construction, a coarse-grained Markov chain never underestimates transitions. Hence a set of states in the Markov chain not having outgoing transitions means that the corresponding Boolean states do not have outgoing transitions. A set of Boolean states with no outgoing transitions must contain at least one attractor. In summary, a coarse-grained Markov chain cannot overcount attractors, i.e., does not predict spurious attractors.

By construction, the SD Markov chain’s recurrent classes are absorbing states (i.e., the minimal trap spaces of the Boolean network). There are two ways in which an SD Markov chain’s absorbing states could undercount attractors. First, attractors may exist outside of minimal trap spaces; this is extremely rare^38^. The second possibility is that a minimal trap space contains multiple attractors. This situation can be ruled out by separately identifying a system’s minimal trap spaces and attractors^39^. We found that in our ensemble of 91 RBNs there was a one-to-one correspondence between minimal trap spaces and attractors.

Even with the one-to-one correspondence, there remains a final source of mismatch between an absorbing state of the SD Markov chain and an attractor of the Boolean network: The absorbing state may map to transient states of the Boolean network that are not part of the attractor and instead are part of its strong basin.

Each absorbing state of the SD Markov chain and each state that belongs to an attractor of the Boolean network can be identified from the infinite (or practically large enough) power of the transition matrix as a state with a non-zero convergence probability to itself. We classify each Boolean state into the four categories of true positive, false positive, true negative, and false negative, based on comparing the Markov chain’s prediction to the ground truth of the Boolean network. These four categories form a confusion matrix (see Methods). For example, false negative means that a Boolean state predicted to be transient by the Markov chain is actually part of an attractor. This can happen in rare cases in principle, but there were no false negatives in our ensemble. Thus every state identified as transient by the SD Markov chain was indeed transient, leading to the recall and the negative predictive value being 1. The average precision of the SD Markov chain is 0.97 (see Table **1**), indicating that only 3.7% of the Boolean states included in an absorbing state of the SD Markov chain are transient states. The worst example of a mismatch is a minimal trap space with 128 states, out of which 96 participate in the attractor (precision of 0.75). Apart from such outliers, precision was very high in most RBNs (it was 1 in 78 out of 91 RBNs). The high average specificity indicates that the SD Markov chain finds 93.4% of transient states. Because the minimal trap spaces of the SD Markov model are shared with the Null and Random Markov chains, these results apply to them as well. To put these results into perspective, we consider a reference confusion matrix that assumes that all states are attractor states. This reference has a recall of 1 and a specificity of 0 by design, meaning that it can find all attractor states and cannot find any transient states. Its precision, which was 0.21, serves as a lower bound for the precision of identifying attractor states. The distributions of the precision and specificity of identifying attractor states, shown in Figure **S1**, indicate that the coarse-grained Markov chains strongly outperform the reference, and have a near-perfect precision and specificity across the RBN ensemble.

In summary, the results of the attractor analysis indicate that the attractors and the minimal trap spaces have a one-to-one relation, and that there are few cases of an attractor not filling a minimal trap space. We conclude that the SD Markov chain predicts attractors and transient states with very high accuracy.

### The SD Markov chain captures basins of attraction

After establishing the almost perfect agreement of the absorbing states of the SD Markov chain with the attractors of the Boolean network, we next evaluate the reachability of attractors, which is embodied by the concept of basin of attraction and by the convergence probabilities. We obtain the convergence probabilities of Boolean states into Boolean attractors using the infinite power of the transition matrix of the Boolean network and use them as the ground truth. The convergence probability into a complex attractor is the sum of the convergence probabilities into the states in the complex attractor. A coarse-grained Markov chain’s transition matrix can be used far more efficiently to obtain the convergence probabilities of the Markov states into absorbing states. These results, when expanded to all the states (as exemplified in Figure **1**), serve as predictions for the convergence probabilities of the Boolean network. To illustrate this evaluation process, Figure **3** shows the convergence probabilities of the Boolean network of Example **1**, and the predictions made by the SD Markov chain and the Random Markov chain defined in Figure **2**.

**Figure 2.**
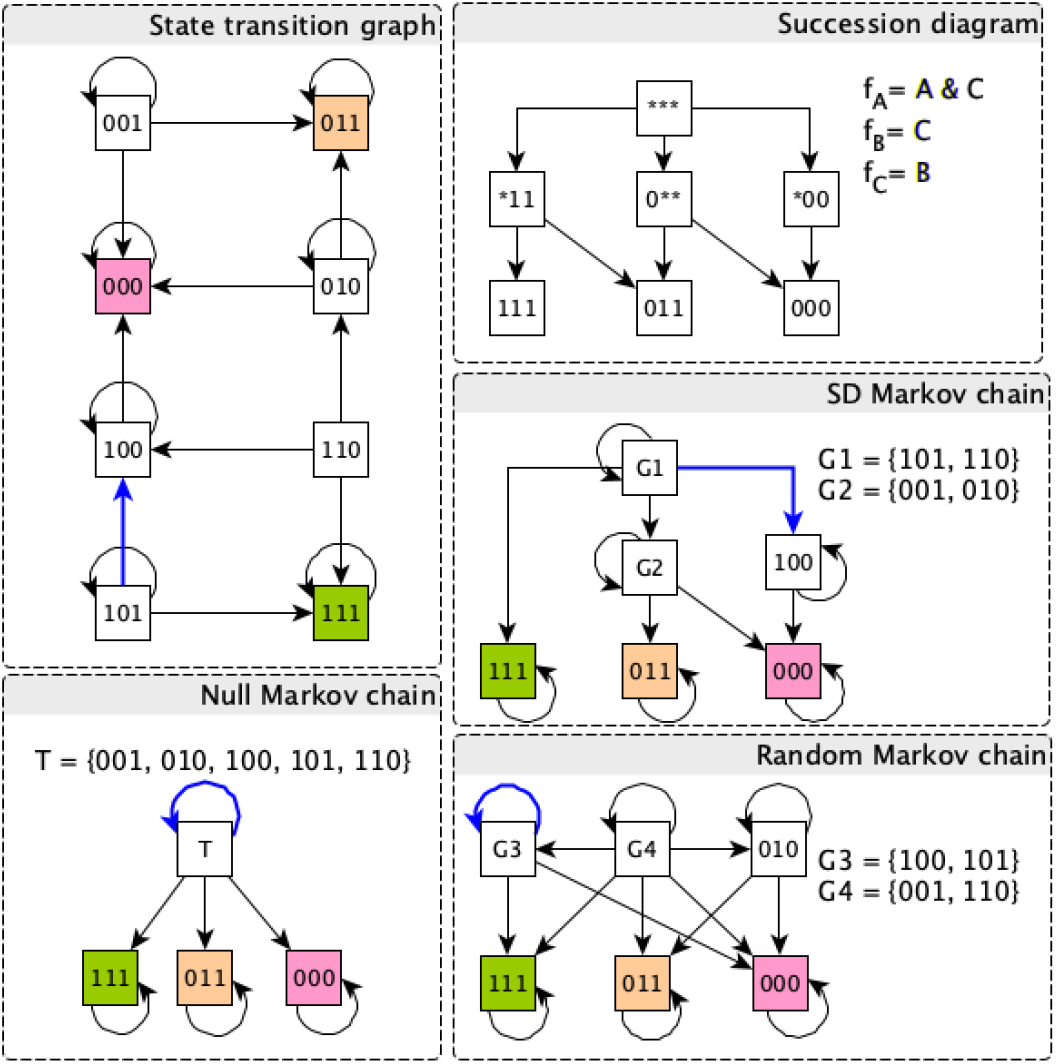
State transition graph and succession diagram of Example **1**, as well as the transition graphs of three coarse-grained Markov chains. The succession diagram contains seven trap spaces. The three minimal trap spaces are point attractors: 111, 011, 000. The trap space *11 contains the states 111 and 011, each of which is already included separately in the minimal trap spaces. The trap space 0** contains four states, two of which (011 and 000) are already separately included. Thus, the group of states exclusive to this trap space is {010, 001}. The trap space *00 contains two states, one of which is 000. The other state, 100, is exclusive to this trap space. The states exclusive to the root trap space are {101, 110}. According to our proposed definition, the SD Markov chain contains 6 states. Two of these are mapped to groups of Boolean states, namely G1= {101, 110} and G2 = {001, 010}. The three minimal trap spaces become absorbing states of the SD Markov chain. The Null Markov chain contains four states, namely the three absorbing states and a state that unites the 5 transient states of the Boolean network. Any Random Markov chain will contain the three absorbing states, as well as three Markov states that together cover the remaining 5 Boolean states. The thick blue line in the state transition graph indicates one example of a decision transition. This decision transition is captured by the SD Markov chain but it falls within group G3 in the Random Markov chain and within group T in the Null Markov chain.

**Figure 3.**
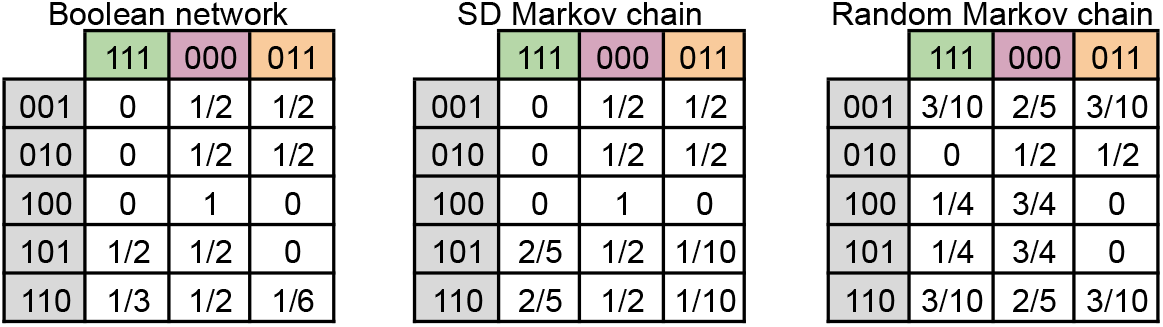
Convergence probabilities of the five transient states of Example **1** into the three attractors (left), compared with the prediction of the SD Markov chain (middle) and the Random Markov chain defined in Figure 2. Strong and weak basins can be read off by counting the number of non-zero entries in each row. For example, the SD Markov chain correctly identifies that 100 is in a strong basin, whereas the Random Markov chain does not. Similarly, attractor reachability can be read off by focusing on the non-zero entries. Although the SD Markov chain correctly identifies that 101 is in a weak basin, it mistakenly predicts that 101 can reach attractor 011.

We evaluated the SD Markov chain’s predictions using three information resolutions within the spectrum from qualitative to quantitative. First, we used the SD Markov chain to classify all Boolean states into states in the strong basin of a single attractor and states that belong to the weak basins of multiple attractors. Second, we used it to classify the reachability between each Boolean state-attractor pair. Third, we evaluated the SD Markov chain-derived convergence probability from each Boolean state to each attractor. For these evaluations, we used the 46 Boolean networks in the RBN ensemble that have more than one attractor.

#### Strong and weak basins

Identifying the states in the strong basin of an attractor is important because these states exhibit commitment to the outcome represented by the attractor, for example, commitment to a specific cell fate. We consider states within an attractor to be in the attractor’s strong basin. When a state is not in a strong basin, it belongs in the weak basins of multiple attractors and converges to each of these attractors with a nonzero probability. For example, based on Figure **3**, in the Boolean network 100 is in the strong basin of the point attractor 000 and 001 is in the weak basins of 000 and 011. If a state of a coarse-grained Markov chain *c*_*j*_ converges into a single absorbing state *a*, the prediction is that all the Boolean states in *G*_*j*_ = π^−1^ (*c*_*j*_) are in the strong basin of the Boolean attractor corresponding to *a*. Due to the overestimation of transitions by coarse-grained Markov chains, this prediction is always correct, thus all positive predictions are true positives. A false negative prediction applies to a Boolean state *s* that maps to a Markov chain state that converges to multiple absorbing states but *s* itself is in the strong basin of an attractor. For example, Figure **3** indicates the Random Markov chain’s prediction that 100 is in the weak basins of 111 and 000, while in reality it is in the strong basin of 000. The definitions of the four classification categories are given in the Methods.

Since false positive predictions are eliminated by design, the precision and specificity of all three coarse-grained Markov chains is 1. As indicated by the violin plots of Figure **4** and the distributions of Figure **S2**, the recall of the SD Markov chain is much higher than the recall of the Random and Null Markov chains over the entire RBN ensemble. The Random Markov chains’ results are nearly identical to those of the Null Markov chains because a strongly connected subgraph of transient states is equivalent to a single transient state in terms of basin inclusion. A characteristic example of an incorrect classification by the Null or Random Markov chains is a Boolean network in which the transient states belong to disjoint strong attractor basins. Grouping any transient states from different attractor basins leads to the erroneous prediction that the states in the group are in a weak basin of attraction.

**Figure 4.**
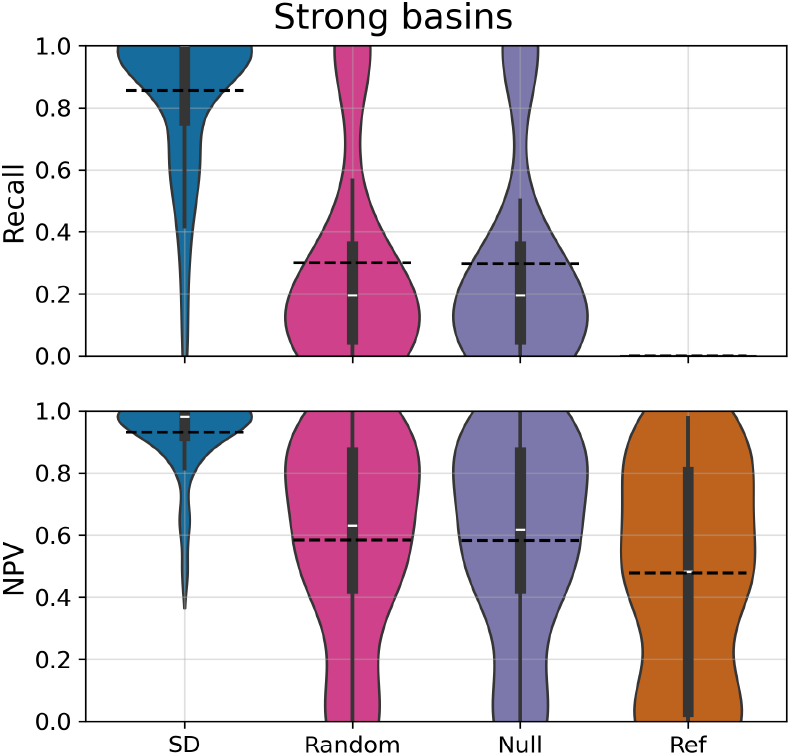
Violin plots of the recall and negative predictive value (NPV) of identifying strong basins by the SD Markov chain (SD), the Random Markov chains (Random), and the Null Markov chain (Null). Also included is a reference that assumes that all states belong to weak basins, and thus has a recall of 0. The median values are indicated with white segments and the averages (also indicated in Table 1) are shown with dashed lines. We use the 46 RBNs that have more than one attractor, to allow for the existence of weak basins. The precision and specificity for all Markov chains is 1 by design. The SD Markov chain’s recall and NPV are concentrated near 1, while the Random Markov chain and Null Markov chain show substantially lower and more dispersed distributions. Two alternative representations of these distributions, shown in Figure S2, confirm that the SD Markov chain exhibits higher recall and NPV compared to the Random and Null Markov chains across the 46 RBNs.

The NPV of the strong basin prediction is akin to the precision of identifying states that belong in the weak basins of multiple attractors. The lower bound for the negative predictive value of a coarse-grained Markov chain is provided by a reference confusion matrix that assumes that any state can reach any attractor. As shown in Figure **4** and Figure **S2**, the SD Markov chain significantly outperforms the Random and Null Markov chains. The latter are somewhat better than the lower bound because they capture the fact that the Boolean states within a minimal trap space are in the strong basin of the attractor corresponding to the trap space.

In summary, these results indicate that the SD Markov chain finds most states that are in a strong basin, and it is fairly precise in predicting that a transient state is in a weak basin.

#### Attractor reachability

To go deeper than identifying strong and weak basins, we evaluate the SD Markov chain by its predictions on the reachability relationship between each Boolean state and each attractor. The construction of a coarse-grained Markov chain ensures that there aren’t any false negative predictions, in which the Markov chain predicts that the Boolean state cannot reach the attractor, but it actually can do so in the Boolean network. Consequently, the recall and NPV of all coarse-grained Markov chains are 1. As indicated in the violin plots of Figure **5** and the distributions of Figure **S3**, the SD Markov chain is very precise in finding the attractors a Boolean state can reach, and identifies most of the attractors that a Boolean state cannot reach. In contrast, the Random and Null Markov chains show only a small improvement compared to a reference confusion matrix that assumes that every Boolean state can reach every attractor. This is because they are similar to the reference in their inability to reflect the cases where a Boolean state mapped to a transient Markov state cannot reach an attractor. They are slightly better than the reference because they capture the fact that the Boolean states within a minimal trap space can only reach the attractor corresponding to the trap space.

**Figure 5.**
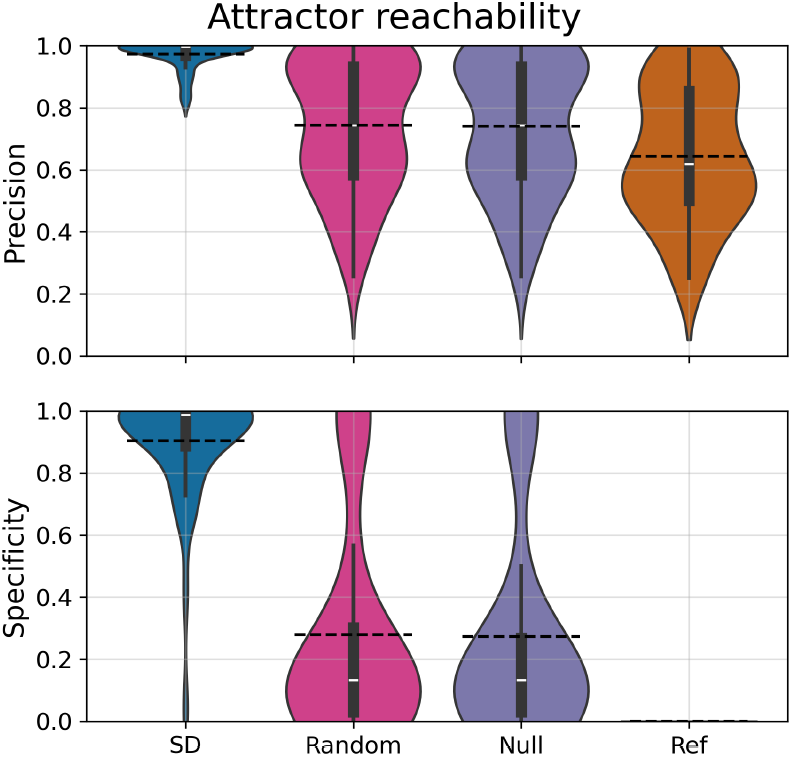
Violin plots of the precision and specificity of identifying reachable attractors from each Boolean state. We use the 46 RBNs that have more than one attractor. The reference assumes that all states reach all attractors, thus its specificity is 0. The average values are shown with dashed lines. The SD Markov chains’ precision and specificity are concentrated near 1. The mass of the Random and Null Markov chains is at lower values, similar to the reference. Alternative representations of this information, indicated in Figure **S3**, confirm that the SD Markov chain outperforms the Random and Null Markov chains over the whole RBN ensemble.

#### Convergence probability into attractors

After evaluating the qualitative reachability between transient states and attractors, we move on to a quantitative evaluation of each Boolean state’s predicted probability of convergence into each attractor. The absorption probability of a state of a coarse-grained Markov chain *c* into an absorbing state *a* will be attributed equally to each transition from a member of π^−1^(*c*) to a member of π^−1^(*a*). Figure **3** illustrates this feature in the case of Example **1**, demonstrating that this equal attribution is an approximation. For example, the SD Markov chain predicts that the convergence probabilities from states 101 and 110 to the attractor 011 are 1/10, 1/10, while the actual convergence probabilities are 0, 1/6. To assess the accuracy of this approximation, we used two quantitative measures of the difference between the Markov-chain-derived convergence probabilities (expanded to all states) and the probabilities in the Boolean network: RMSD and KLD. We determined the RMSD and KLD for each Boolean state (i.e., each row of the convergence matrix), then averaged over states.

To put these results into perspective, we determined the RMSD and KLD of a reference that assumes equal convergence to each attractor. Note that this reference does not constitute a limiting case. As shown in Figure **6** and Figure **S4**, both measures indicate that the SD Markov chain is much better at predicting the convergence probability into attractors than the Random and Null Markov chains, which in turn are better than the reference. Notably, the average RMSD value of 0.01 for the SD Markov chain means that it can predict the convergence probabilities of every state with an around 1 percentage point error.

**Figure 6.**
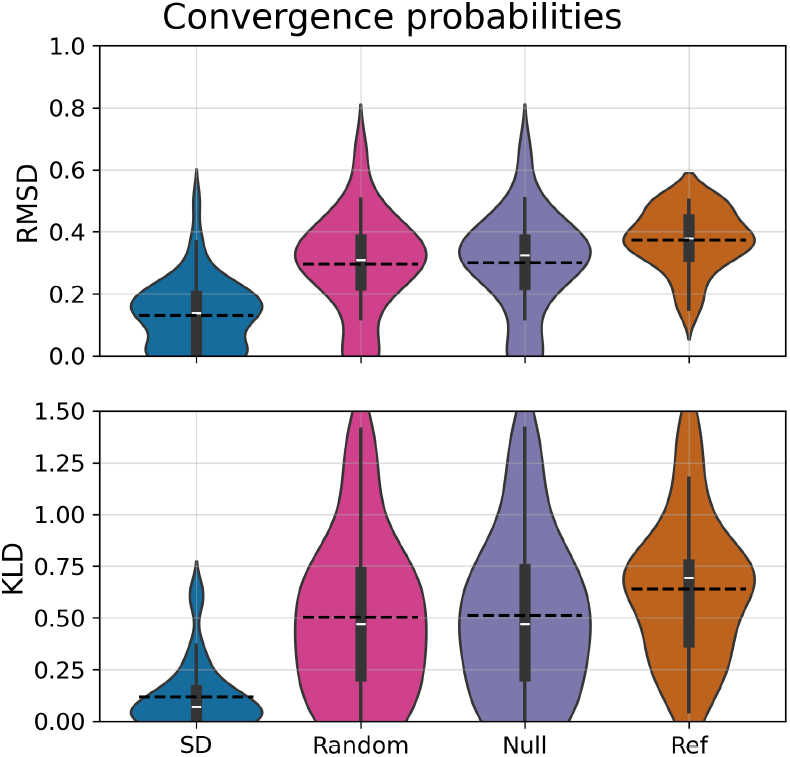
Violin plots of the RMSD and KLD between the convergence probabilities of the Boolean network and the coarse-grained Markov chains. The median values are indicated with white segments and the averages (also indicated in Table 1) are shown with dashed lines. We use the 46 RBNs that have more than one attractor. The reference assumes convergence to all attractors with equal probability. The SD Markov chain’s RMSD and KLD are concentrated at low values. The mass of the Random and Null Markov chains is at higher values, similar to the reference. Alternative visualizations of the distributions, shown in Figure **S4**, confirm that the SD Markov chain consistently outperforms the Random and Null Markov chains and the reference.

### The SD Markov chain captures basin fractions and average node values

The SD Markov chain provides much more in-depth predictions regarding the Boolean network’s attractor basins than simulation methods. It is still informative to evaluate its predictions for measures that are usually estimated with simulations that sample random initial conditions, namely the basin fractions of attractors and average node values. We calculated these metrics by taking appropriate weighted averages over convergence probabilities.

#### Basin fractions

In a Boolean network with multiple attractors, the basin fraction of an attractor indicates the probability of converging into it from an initial state selected uniformly randomly (see Methods). The biological interpretation of a basin fraction could be, for example, the fraction of one of multiple differentiated cell types obtained from a mixture of progenitor cells. We calculate the basin fractions using the convergence probability of each state. For example, the basin fractions of Example **1** are 0.5, 0.27, 0.23 for the point attractors 000, 011, 111, respectively. The predicted basin fractions of Example **1** according to the SD Markov chain are 0.5, 0.275, 0.225, which closely matches the actual basin fractions.

For each member of the RBN ensemble, we determined the basin fractions of the absorbing states of the SD Markov chain and quantified their difference from the basin fractions of the Boolean network’s attractors using the RMSD and KLD. Both of these metrics are applied on a vector whose length is the number of Boolean attractors. We used a reference that assumes equal convergence to each attractor, which thus implies an equal basin fraction. We found that the SD Markov chain is significantly more accurate at capturing the Boolean basin fractions than the Random or Null Markov chains or the reference (see Figure **7** and Figure **S5**). The average RMSD value of 0.05 for the SD Markov chain means that it can predict the basin fractions with an around 5 percentage point error.

**Figure 7.**
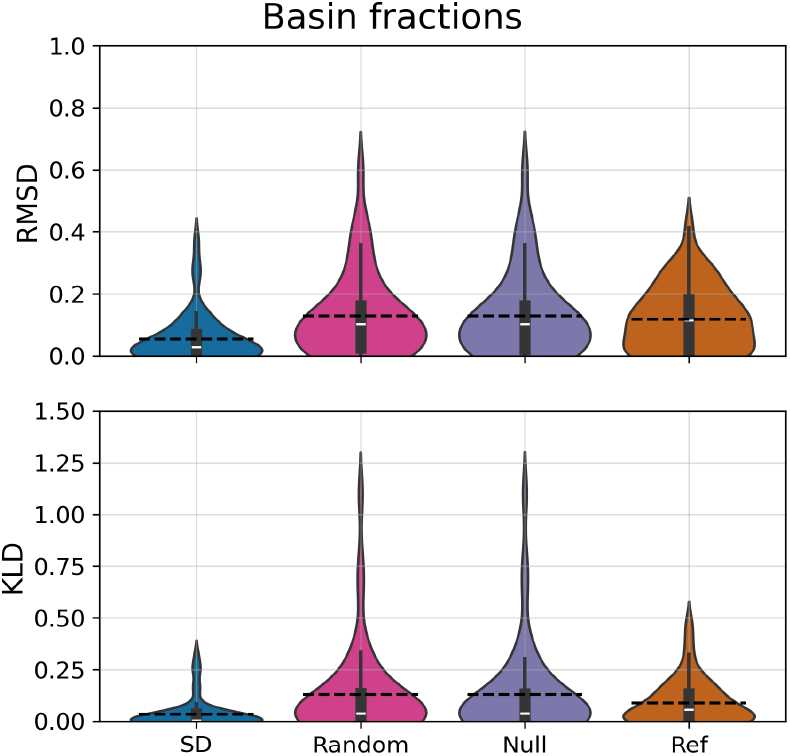
Violin plots of the RMSD and KLD between the basin fractions of the attractors of the Boolean network and the corresponding absorbing states of the coarse-grained Markov chains. We use the 46 RBNs that have more than one attractor. The reference assumes convergence to all attractors with equal probability. Alternative representations of this information, indicated in Figure **S5**, confirm that the SD Markov chain outperforms the Random and Null distributions and the reference over the whole RBN ensemble.

#### Average node values

The average value of each node of the Boolean network can be interpreted as the average expression or activation level of the respective element in experimental assays that measure cell populations. This average value can be calculated using the value of the node in each attractor and the convergence probability to each attractor (see Methods for the formula). For the Boolean network in Example **1**, the average value of node **A** is 0.229 and the average value of nodes **B** and **C** is 0.5. The SD Markov chain predicts an average value of 0.233 for node **A** and 0.5 for nodes **B** and **C**.

We determined the average node values for each Random Boolean network with more than one attractor. As indicated in Table **1** and Figure **8**, the RMSD of the SD Markov chain is 0.05, which indicates that the SD Markov chain can determine the average node values with an average error of 0.05. This is much lower than the RMSD of the Random and Null Markov chains, which in turn is much lower than the RMSD of a reference that assumes an average node value of 0.5 for all nodes. The violin plots of Figure **6** and the distributions of Figure **S6** indicate that the SD Markov chain’s RMSD and KLD values are consistently lower than those of the Random and Null Markov chains and of the reference.

**Figure 8.**
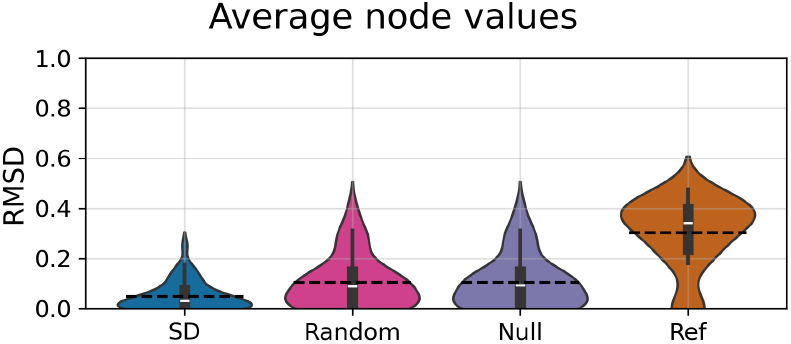
Violin plots of the RMSD obtained from the average node values of the Boolean network and the coarse-grained Markov chains. The plot shows results for 46 RBNs that had more than one attractor. The reference assumes a value of 0.5 for every node. Alternative representations of this information are available in Figure **S6**.

### The SD Markov chain captures decision transitions

Having established the good accuracy of the SD Markov chain in reflecting the basins of attraction of the Boolean network, we evaluate its performance in identifying basin boundaries. In particular, we introduce the concept of decision transition by adapting the concept of basin boundary defined by Bérenguier and colleagues^35^. Intuitively, a decision transition is a transition along which there is a decrease in attractor reachability, e.g., from a weak basin to a strong basin. We provide a more detailed definition in the Methods. In the Boolean network of Example **1** almost all state transitions are decision transitions. The only exception is 100 → 000, because both states are in the strong basin of 000. The decision transition 101 → 100 is indicated in blue in Figure **2**.

A coarse-grained Markov chain will reflect decision transitions as pairs of Markov states *c*_*j*_, *c*_*k*_ such that (i) there is a *c*_*j*_ → *c*_*k*_ transition, (ii) *c*_*j*_ converges to multiple absorbing states, and (iii) *c*_*k*_ converges to a strict subset of the absorbing states to which *c*_*j*_ converges. Since the definition of the coarse-grained Markov chain loses information on the differences between the Boolean states that are in *G* = π^−1^ (*c*) = {*s* ∈ *S*|π(*s*) = *c*} grouped into a state of the Markov chain, it is not possible to make a prediction on which specific transition from a state *s*_*j*_ ∈ *G*_*j*_ to a state *s*_*k*_ ∈ *G*_*k*_ exists in the Boolean network. Thus, we classify every transition *s*_*j*_ → *s*_*k*_ that exists in the Boolean network according to the prediction arising from the *c*_*j*_ → *c*_*k*_ transition of the SD Markov chain (see Methods). An example of a false negative prediction is a decision transition of the Boolean network that falls between Boolean states corresponding to the same Markov state *c* (i.e., a decision transition *s* → *s*’ where *s, s*’ ∈ π^−1^ (*c*)). This is the case for the Random Markov chain illustrated in Figure **1**: the decision transition 101 → 100 is inside the group G3={100, 101}.

We determined the precision, recall, NPV, and specificity of identifying decision transitions for the 46 Boolean networks in the RBN ensemble that have more than one attractor). Twelve models did not have any decision transitions, which the SD Markov chain correctly identified (and had undefined precision). We find that the SD Markov chain captures decision transitions to a remarkably good extent (see Table 1 and the distributions in Figure **S7**). The high average precision of the SD Markov chain, 0.91, indicates that if *c*_*j*_ → *c*_*k*_ is a decision transition in the Markov chain, the transitions *s*_*j*_ → *s*_*k*_ indeed are decision transitions in most cases. The good average recall of the SD Markov chain, 0.82, indicates that it misses relatively few decision transitions. The Null and Random Markov chains’ precision and recall were much lower than those of the SD Markov chain due to a significant number of false positives (see Figure **S7**). For example, these Markov chains predicted decision transitions in 6 Boolean networks that had none by grouping multiple transient states that belong in the strong basin of different attractors. The SD Markov chain has almost perfect NPV and specificity, meaning that it identifies non-decision transitions with very high precision and recall.

### The SD Markov chain captures sequences of events

The results presented in the previous sections show that the SD Markov chain successfully reflects the long-term behaviors of the Boolean network and the decisions that lead to them. A remaining question whose answer would be insightful is: how many different sequences of events (i.e., trajectories) exist, and how likely are they? A sequence of events could correspond to the sequential activation of nodes in a signal transduction pathway. Alternative sequences of events leading to the same outcome could correspond to alternative pathways eliciting the same cell state. Here we show that the SD Markov chain reflects the trajectories in the Boolean network (i.e., the paths in its state transition graph), and accurately summarizes alternative sequences of events and their probabilities.

#### Existence of trajectories

The coarse-grained Markov chain overestimates transitions by design. This means that the lack of transitions from *c*_1_ to *c*_2_ in a coarse-grained Markov chain indicates that no transitions exist in the Boolean network from any Boolean state *s*_1_ ∈ *G*_1_ = π^−1^(*c*_1_) to any *s*_2_ ∈ *G*_2_. Furthermore, if a path exists in the STG of a Boolean network from state *s*_1_ to *s*_2_, the corresponding path from *c*_1_ to *c*_2_ exists in the Markov chain. Unfortunately, the inverse is not true in general. When the Markov chain’s state transition graph contains a two-transition path *c*_1_ → *c*_2_ → *c*_3_, this indicates that there exists a pair of Boolean states *s*_1_ and *s*_2_ connected by a transition, and another pair of Boolean states *s*_2_’ ∈ *G*_2_ and *s*_3_ ∈ *G*_3_ connected by a transition. However, there is no guarantee that *s*_2_’ is reachable from *s*_2_, and thus the existence of a path in the Markov chain does not imply the existence of a corresponding path in the Boolean network’s STG. Indeed, in the Random Markov chain of Example **1**, the *G*4 → *G*3 → 111 path does not have a corresponding path in the STG.

Thus the question is: if a path exists in the Markov chain, how likely is it for that path to exist in the STG of the Boolean network? This question is meaningful if the paths in the Markov chain can distinguish among the possible sequences of events in the Boolean network. The Null Markov chain only has paths of length 1, namely edges from the transient group to the attractor groups, and hence cannot reflect any sequences of events. The Random Markov chain forms an almost complete graph of groups of transient states, predicting that every path is possible, which doesn’t distinguish among sequences of events. Therefore, we performed this analysis only for the SD Markov chain, which has a sparse, acyclic structure of state transitions.

We evaluated the correspondence of paths in the SD Markov chain with trajectories in the Boolean network using the 68 Boolean networks in the RBN ensemble whose SD Markov chains have paths of length 2 or longer. For every such path, *c*_1_ → *c*_2_ → … → *c*_*n*_ in the SD Markov chain, we checked whether a trajectory exists in the Boolean network that starts from a Boolean state in *G*_1_ = π^−1^(*c*_1_), goes through states in *G*_1_, *G*_2_,.. in a sequence, and arrives to a state in *G*_*n*_ = π^−1^(*c*_*n*_). If the answer is yes, the SD Markov chain’s path is classified as true positive (TP). Otherwise, it is classified as false positive (FP). There are no false negatives by design, thus the recall of the SD Markov chain is 1. The precision was 1 for 67 RBNs and 0.98 in a single RBN. This latter RBN stands out in having the largest number of attractors (6) and the second largest number of Markov states and transitions (30 and 80, respectively). Thus, it is not surprising that there exist 7 paths of the SD Markov chain (out of 312) that do not have a corresponding path in the original system’s STG.

The result shows that the SD Markov chain provides an efficient summary of all trajectories of the Boolean network. This is likely due to the efficiency with which the SD-based groups partition the state space and reflect the decrease in the freedom of the system. Also note that the acyclic nature of the succession diagram helps in enumerating all paths.

#### Trajectory probabilities

Having established the good correspondence between the existence of paths, we investigated the path probabilities. Our chosen method was first-step analysis^23^, which is a way to determine the probability *P*(*x* → *y*) of a Markov chain eventually converging to a state *y* when starting from a state *x* by following the transitions allowed from state *x* (see Methods). First-step analysis of Example **1**, described in the Methods, indicates that the probability of the *G*1 → *G*2 → 000 path is 0.10 in the SD Markov chain. First step analysis can also be used to determine the aggregated probability of Boolean trajectories that correspond to a path of the SD Markov chain by solving a system of linear equations (see Methods). First-step analysis indicates that the probability of the corresponding trajectories {101, 110} → {001, 010} → 000, when assuming equal probability of the two initial states, is 0.08, a close but not perfect agreement.

We determined the RMSD between the convergence probabilities over paths of length 2 or longer in the SD Markov chain and the corresponding convergence probabilities in the Boolean network. As a reference, we randomized the nonzero probabilities in the SD Markov chain. Note that this reference is as effective as the SD Markov chain in reflecting the existence of trajectories of the Boolean network. The average RMSD for the SD Markov chain is much lower than that of the reference (see Table **1**), and the cumulative distribution of the RMSD of the SD Markov chain lies consistently above that of the reference (see Figure **S8**) indicating the superior accuracy of the SD Markov chain.

The success of the SD Markov chain in reflecting the trajectories of the Boolean network allows its use to identify the independent sequences of events that jointly determine the convergence probability into attractors. In the case of Example **1**, the SD Markov chain indicates that there are two paths that start from *G*1 and converge into 000: one through *G*2 (i.e., through the trap space 0**) and another through 100. The SD Markov chain indicates that the probability of convergence to 000 through 100 is fourfold higher (in the Boolean network the ratio is 5). This result is non-trivial and actionable. For example, if one wanted to prevent the 110 → 000 convergence and could eliminate only one of these two sequences of events, this result indicates that the 110 → 100 → 000 path should be prioritized for elimination.

## Discussion

Understanding a complex biological system made up of many non-identical elements through dynamic modeling is hindered by the high dimensionality of the model’s state space. Here, we present progress toward addressing this challenge in the context of stochastic asynchronous Boolean models of biological systems. We show that the succession-diagram-based Markov chain achieves a biologically interpretable coarse-graining by grouping Boolean states that share causal constraints defined by trap spaces. The SD Markov chain retains the explanatory strength of the succession diagram in describing the decisions that channel the dynamics, and goes significantly beyond it by describing the probability of each decision. We find that the SD Markov chain successfully captures the attractor landscape of the full Boolean model, including the composition of attractors, the basins of attraction of each attractor, and the boundaries between basins. The paths of the SD Markov chain reflect the trajectories of the Boolean model. The convergence probability along these paths is indicative of the probability of sequences of events in the Boolean model.

The attractors of biomolecular systems correspond to stable cell fates or phenotypic states, and the transitions among them represent differentiation or reprogramming^46^. The influence of stochasticity and noise on the transitions among phenotypic states has long been recognized^47,48^. Markov chain modeling is a powerful framework for incorporating stochasticity. Given a state transition matrix of a tractable size, Markov chain analysis can determine the convergence probabilities into attractors. De novo construction of a Markov model of a biological system necessitates prior knowledge of a small repertoire of states and of the transition probabilities among them. Markov models have been successfully constructed for certain systems, such as to describe the transitions among conformations of biomolecules^42^. Yet, cell states emerge from the interactions of many and diverse biomolecules. Understanding the transitions among cell states necessitates a description of the dynamics of the biomolecular system that underlies these cell states, for example via a Boolean model. We envision a two-step modeling process to describe transitions among cell states: First, incorporate the biomolecular interactions and regulatory mechanisms into a Boolean model that reflects the stochasticity in the system. This model reveals the emergent cell states and the decisions that lead to each. Next, use the succession diagram of the Boolean model to construct the SD Markov chain. The dramatic reduction of the number of states in the SD Markov chain compared to the number of states of the Boolean system allows the construction and analysis of the SD Markov chain’s transition matrix, revealing the sequences of events and the probabilities corresponding to each transition among cell states. This synergistic combination of the strengths of the Boolean and Markov modeling frameworks is made possible by the coarse-graining method proposed here.

Our current work focused on evaluating the SD Markov chain in cases where it could be constructed from the STGs of small Boolean networks. The success of this evaluation opens the way for application of the SD Markov chain for Boolean networks for which it is not possible to obtain or handle the state transition graph. Transition probabilities among each pair of state groups can be determined by single-step simulations that start from the states in the first group. The optimal implementation of these simulations will be the topic of future work.

The SD Markov chain enables new ways to analyze cell fate plasticity and control strategies in high-dimensional biomolecular networks. For example, the SD Markov chain can be used to fine-tune control strategies that rely on the succession diagram^31,49,50^. Because SD Markov chain analysis reflects the trajectories of the system and their probabilities, it can identify the interventions and their durations needed for any sequence of events to happen.

As the SD Markov chain derived from a Boolean model is a model in its own right, it can be refined to serve as a better descriptor of the system than the original model. This model refinement is made possible by the manageable size of the state transition matrix. For example, the transition probabilities can be adjusted, as long as the transition matrix remains stochastic. Such adjustments will allow the study of the manner in which the distribution of attractor basins becomes more even, or more uneven. Adjusting the model to fit known observations will enable more precise predictions for settings that were not studied before.

Another way of refining the SD Markov model is by incorporating multiple sources of stochasticity and varying their strength. The variability of the timing of biomolecular processes has been incorporated into Boolean models by proposing multiple update schemes^2^. The stochasticity that impacts regulation has been modeled by introducing probabilistic update functions^51,52^. These stochastic models were so far only amenable to simulation studies that sample the state space. Now these sources of stochasticity can be incorporated into the SD Markov chain’s state transition matrix. The states of the Markov chain would be the same, the only change would be to the transition probabilities. The tractable size of the SD Markov chain’s transition matrix allows full analysis of the SD Markov chain. The results can then be translated back to the original system.

In particular, it is straightforward to determine the SD Markov chain for other update schemes. The Markov chain’s correct reflection of prohibited transitions originates in the succession diagram’s structure and is independent of the update scheme. The more transitions allowed by the update scheme, the smaller the loss of information due to the Markov chain’s overestimation of transitions. Synchronous update is handled poorly due to its deterministic nature. Update schemes with more transitions than our current implementation, such as the most permissive update^53^, would have a decreased likelihood of mismatches between minimal trap spaces and attractors, and increased agreement between the SD Markov chain and the Boolean network in attractor reachability and the existence of paths.

Probabilistic update functions, such as those used in Boolean networks with perturbations^51^ and models with stochastic propensity^52^, can also be incorporated into the transition matrix of the SD Markov chain. Analysis of the SD Markov chain in case of an increasing degree of stochasticity will allow the tracing of the gradual destabilization of attractors.

Logical models that contain nodes with multiple discrete levels have been fruitfully used to describe biological systems^3,54^. The trap spaces and succession diagram of multi-valued logical models can be determined the same way as those of Boolean models^55^. As these models have an even larger state space, the reduction in the number of states offered by the SD Markov chain will be even more welcome. The SD Markov models derived from multi-level discrete models may reveal more subtle effects of timing variability and stochasticity on the attractor landscape.

Boolean modeling followed by SD Markov chain analysis provides a discrete analogue of the continuous-variable stochastic methods used to describe the probabilistic attractor landscape of cell differentiation^56,57^. It is a complementary approach that is especially suited for systems with several attractors. Future research may be able to combine the strengths of the succession diagram and of stochastic continuous models. Indeed, continuous models within a broad class relevant to biomolecular systems^58^ exhibit identifiable trap spaces (positive-invariant sets)^59^, opening the possibility of constructing a succession diagram. We expect that the SD Markov chain will become a practical and effective reduced model for multiple type of dynamic models. More generally, the SD Markov chain’s bridging of qualitative and quantitative modeling establishes it as a scalable framework for exploring how structure constrains dynamics in complex biological networks.

## Methods

### Choosing trap spaces of the succession diagram as states of the Markov chain

The states of the SD Markov chain are trap spaces of the Boolean model. We follow a selection process to ensure that each state of the Boolean model is assigned to a single state in the Markov chain.

We first build a pool of trap spaces. Each node of the succession diagram is a trap space, which we include in the pool of trap spaces. The edges of the succession diagram represent the stable motifs^29,37^. Each of these edges, combined with its starting node, also defines a trap space, which we also include in the pool. We assign a state of the Boolean model to the smallest possible trap space it belongs to. For example, a state in a minimal trap space is assigned to the minimal trap space, and not to the larger trap spaces that contain the minimal trap space. In rare cases, there is ambiguity in the assignment. This happens when the state belongs to multiple edges of the succession diagram, but not to any smaller trap spaces. In such cases, we add the intersection of the trap spaces (which is also a trap space) into the pool, then the state belongs to this trap space. This assignment process ensures that each state of the Boolean model corresponds to a single state in the Markov chain.

### Random Boolean Network (RBN) ensemble

We followed the NK model proposed by Stuart Kauffman^60^, in which each of the *N* nodes receives *K* edges from randomly selected nodes, and the update functions are chosen such that each of the 2 ^*K*^ input combinations yields an output of 1 with probability *p*. We generated an ensemble of 100 Boolean networks using *N* = 10, *K* = 3 and *p* = 0. 21, which ensure a good variety of dynamic behavior. As in some members of the ensemble there were nodes whose Boolean function was a constant, we performed percolation of constants, and kept the non-constant nodes only. This caused the total reduction of 9 Boolean networks, leaving 91 systems with 1 to 10 nodes, with a median of *N* = 8. Sixteen members of the RBN ensemble do not have any transient states. Forty-five members of the RBN ensemble have a single minimal trap space; the rest have multiple minimal trap spaces.

### Biodivine Boolean Models (BBM) ensemble

The BBM ensemble^32^ includes 212 published Boolean models of biological systems. We used the results of the succession diagram identification analysis of this model ensemble with the **biobalm** tool^31^ to compare the size of the succession diagram with the size of the state space. We considered the analysis in which the input nodes are fixed in the state 1. To account for the fixed inputs, we did not count the input nodes when determining the size of the state space of each model. The number of variable (non-input) nodes in the BBM ensemble ranges from 4 to 302, with a median of 30. There are 129 models with more than 20 nodes, which therefore have more than 1 million states. The succession diagram of four models could not be determined in 10 minutes, and six additional models had more than 600 trap spaces in the succession diagram. The median size of the succession diagram was 3, and 183 models had 20 or fewer trap spaces in the succession diagram.

### Comparing the SD Markov chain, Null Markov chain and Random Markov chains

The SD Markov chain’s states are the trap spaces of the succession diagram of the Boolean model. The Null Markov chain has the same absorbing states as the SD Markov chain, and merges all transient Boolean states into a single Markov state. The Random Markov chains have the same number of states as the SD Markov chain, and map Boolean states to Markov states uniformly randomly. Each Boolean network that doesn’t have any transient states maps to a Markov chain that is composed of absorbing states only. The three coarse-grained Markov chains share absorbing states, so they are identical for such Boolean networks. Any Boolean network whose transient states fall into the same trap space will also lead to identical coarse-grained Markov chains. We compared the three Markov chains for the the 71 RBNs in which the three Markov chains are not identical.

The maximal number of transitions in a Markov chain with *N*_*t*_ transient states and *N*_*a*_ absorbing states is *N*_*t*_ · *N*_*t*_ + *N*_*t*_ · *N*_*a*_ + *N*_*a*_. Here, the first term indicates a complete subgraph of transient states, the second term indicates transitions from every transient state to every absorbing state, and the third term indicates the self-edges of the absorbing states. The Null Markov chain usually has maximal connectivity, the rare cases in which there is an isolated absorbing state being exceptions. Since the Null Markov chain has a different number of states than the SD and Random Markov chains, their densities cannot be directly compared. As described earlier, we find that the SD Markov chains have around 50% of the maximal connectivity, while the Random Markov chains have near-maximal connectivity.

### Statistical classification using a confusion matrix

These classifications are used to compare a feature of a coarse-grained Markov chain with the corresponding feature in the Boolean network’s state transition graph. Each occurrence of this feature is classified as a true positive (TP) when the feature exists in both systems, false positive (FP) when the feature exists in the Markov chain but not in the Boolean network, false negative (FN) when the feature does not exist in the Markov chain but it does exist in the Boolean network, and true negative (TN) when the feature does not exist in either system.

It is important to note that the number of actual positives and actual negatives are fixed by the Boolean network of interest, and determine the sums TP + FN and FP + TN, respectively. The Markov chain produced from the Boolean network determines the distribution of the two respective classifications in the relevant category. For example, for a specific Markov chain 20% of the actual positives is classified as TP and 80% is classified as FN. In a lot of cases, the Markov chain we construct has a FN (or FP) value of 0 by design; then the quality of the Markov chain is determined by its ability to correctly identify the negatives (or positives).

Precision, defined as TP/(TP+FP), indicates the fidelity of the Markov chain in predicting the occurrence of the feature. Recall, defined as TP/(TP+FN), shows the Markov chain’s ability to identify all occurrences of the feature. Specificity, defined as TN/(TN+FP), shows the extent to which the Markov chain identifies the cases in which this feature does not exist. The negative predictive value (NPV), defined as TN/(TN+FN), indicates the fidelity of the Markov chain in identifying the cases in which the feature does not exist. Throughout our analysis, we find these values for each Markov chain on each Boolean network, and average them over the ensemble of Boolean networks.

### Specific assignments of TP, FP, FN and TN in the context of attractors and their reachability

When evaluating the capacity of the SD Markov chain to capture Boolean attractors, a Boolean state is a true positive (TP) if it maps to an absorbing state of the SD Markov chain and is part of an attractor of the Boolean network. A Boolean state is a false positive (FP) if it maps to an absorbing state of the SD Markov chain but is not part of an Boolean attractor. The Boolean state can be a false positive when it is a transient state that lies within a minimal trap space. A state that is transient according to both the SD Markov chain and the Boolean network is a true negative (TN). A Boolean state would be a false negative (FN) if it did not map to an absorbing state of the SD Markov chain but it would be a part of an attractor of the Boolean network. Such false negatives are impossible due to the construction of the SD Markov chain.

When evaluating strong basins, each state of the Boolean network that (i) belongs to a state of the SD Markov chain that converges to a single absorbing state and (ii) is in the strong basin of the Boolean attractor corresponding to that absorbing Markov state is classified as a true positive (TP). A false positive (FP) would be a Boolean state which is predicted to be in a strong basin, but in reality it is in a weak basin. This is not possible due to the overestimation of transitions by coarse-grained Markov chains. A Boolean state is classified as a true negative (TN) if it corresponds to a Markov state that can converge to multiple absorbing states and indeed it is in the weak basin of multiple attractors. A false negative (FN) is a Boolean state that corresponds to a Markov state that converges to multiple absorbing states but the state itself is in the strong basin of an attractor. A reference confusion matrix that assumes that all states are in weak basins has TP=FP=0. The Null and Random Markov chains have a nonzero value of TP, because they capture the fact that the Boolean states within a minimal trap space are in the strong basin of the attractor corresponding to the trap space.

When evaluating attractor reachability, if the Boolean state can reach the attractor both according to the Markov chain and in the Boolean network, the relationship is classified as a true positive (TP). If the Markov chain predicts that the Boolean state can reach the attractor but the state is not part of the attractor basin in the Boolean network, the relationship is classified as a false positive (FP). An example of a FP in Table **3** is the SD Markov chain’s prediction that 101 can reach 011. True negatives (TN) are defined similarly to TP. A false negative (FN) would be a case in which the Markov chain predicts that the state cannot reach the attractor, but it can do so in the Boolean network. By construction of the coarse-grained Markov chain, false negatives are not possible. A reference confusion matrix that assumes that all states can reach all attractors has TN=FN=0. The Null and Random Markov chains have a nonzero value of TN, because they capture the fact that the Boolean states within a minimal trap space can only reach the attractor corresponding to the trap space. Nevertheless, their TN value is small, because they rarely capture cases in which a Boolean state mapped to a transient Markov state cannot reach an attractor.

### Quantifying the information loss of probabilities

We obtain various probabilities that represent the dynamics of the system, such as the convergence probability to each attractor. To compare the prediction gained from the coarse-grained Markov chain to the ground truth obtained from the Boolean network, we use two measures: root-mean-square difference (RMSD) and Kullback-Leibler divergence (KLD).

RMSD can be roughly interpreted as the error for each predicted value. In case of perfect agreement, the RMSD is 0. Larger errors have a disproportionately large effect on RMSD. Consequently, the RMSD is sensitive to outliers. We use RMSD to evaluate predictions for probability distributions and average node value predictions.

KLD is frequently used to measure the difference between a model probability distribution *Q*(*x*) and a real probability distribution *P*(*x*). The KLD is defined as 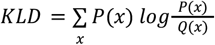, We apply this measure to two probability distributions: i) the convergence probability, which for each initial state describes the probability to reach each attractor, and ii) the basin fraction, which is the probability that a randomly selected state will converge to a given attractor. In both cases, the Markov-chain-derived convergence probabilities serve as the model *Q*(*x*), and the Boolean network’s convergence probabilities serve as the real distribution *P*(*x*). Note that KLD diverges if *Q*(*x*) = 0 for any *x* for which *P*(*x*) ≠ 0. In other words, the loss of information is considered infinite if an event that is possible in the real system is considered impossible in the model prediction.

### Calculating basin fractions and average node values

Consider a Boolean network with *N* nodes. The basin fraction of attractor ***A*** is

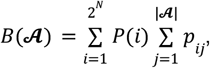

where *P*(*i*) is the probability distribution of the initial states *i, p*_*ij*_ is the convergence probability from the initial state *i* to the attractor state *j*, and |***A***| indicates the number of states in attractor ***A***.

The average value of each node can be calculated as

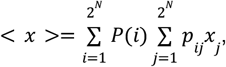

where *P*(*i*) is the probability distribution of initial states *i, p*_*ij*_ is the convergence probability from state *i* to state *j* (which will be 0 if state *j* is not part of an attractor), and *x*_*j*_ is the value of the node in state *j*.

In our work, we assume that each state has an equal chance of being an initial state, namely 2^−*N*^ for both cases.

When calculating the basin fractions and average node values predicted by a coarse-grained Markov chain, the convergence probability is derived from the infinite power of the Markov chain’s transition matrix, expanded to all nodes of the Boolean network. The basin fractions are calculated for the set of Boolean states mapped to each absorbing state of the Markov chain.

### Defining and classifying decision transitions

Consider Boolean states *s*_1_ and *s*_2_ such that (i) there is an *s*_1_ → *s*_2_ transition, and (ii) both states can reach attractor ***A***_2_ but only *s*_1_ can reach attractor ***A***_1_. Then the *s*_1_ → *s*_2_ transition belongs to the boundary of the basin of attraction of ***A***_1_. Indeed, if all such boundary transitions were removed, the remaining basin of attraction of ***A***_1_ will be isolated from the rest of the basins. Since the *s*_1_ → *s*_2_ transition represents a transition from a less restricted basin (the union of the basins of attractors ***A***_1_ and ***A***_2_) to a more restricted basin (the basin of ***A***_2_), one can also view this transition as a decision. We will call such transitions “decision transitions”. A decision transition for a coarse-grained Markov chain is defined similarly, as explained earlier.

Since a coarse-grained Markov chain loses information on the differences between the Boolean network’s states that map to the same state of the Markov chain, we classify every *s*_*j*_ → *s*_*k*_ transition that exists in the Boolean network according to the prediction arising from the *c*_*j*_ = π(*s*_*j*_) → *c* _*k*_ = π(*s*_*k*_) transition of the SD Markov chain. We define a true positive (TP) as a transition *s*_*j*_ → *s*_*k*_ that is a decision transition in the Boolean network with the property that *c*_*j*_ → *c*_*k*_ is a decision transition in the SD Markov chain. A false positive (FP) is a case in which *s*_*j*_ → *s*_*k*_ is not a decision transition but *c*_*j*_ → *c*_*k*_ is a decision transition in the SD Markov chain. This happens when some other state *s*_*j*_ ′ ∈ π^−1^(*c*_*j*_) can reach attractors that *s*_*j*_ cannot reach, but *s*_*j*_ and *s*_*k*_ can reach the same attractors. As a result, *c*_*j*_ overestimates the reach of *s*_*j*_.

### First-step analysis of Markov chains

First-step analysis^23^ is a way to determine the conditional probability of a Markov chain eventually converging to a state *y* when starting from a state *x, P*(*x* → *y*), by following the transitions allowed from state *x*. Consider that state *x* can stay unchanged with probability *p*_*xx*_, transition into state *z* with probability *p*_*xz*_, or transition into state *t* with probability *p*_*xt*_.

First step analysis yields *P*(*x* → *y*) = *p*_*xx*_ *P*(*x* → *y*) + *p*_*xz*_ *P*(*z* → *y*) + *p*_*xt*_ *P*(*t* → *y*).

The same analysis is then applied to formulate equations for *P*(*z* → *y*) and *P*(*t* → *y*), continuing until all paths to *y* are covered. This recursive process yields a system of linear equations, as many equations as many states can reach *y*.

It is also possible to restrict the paths included in the analysis and calculate, e.g., the conditional probability of converging to *y* following only paths through *z*. Let’s assume that state *z* can stay unchanged or transition to *y*. Then *P*(*z* → *y*) = *p*_*zz*_ *P*(*z* → *y*) + *p*_*zy*_, which yields *P*(*z* → *y*) = *p*_*zy*_ /(1 −*p*_*zz*_). Then, the conditional probability of the *x* → *z* → *y* path satisfies *P* (*x* → *z* → *y*) = *p*_*xx*_ *P*(*x* → *z* → *y*) + *p*_*xz*_ *p* _*zy*_ /(1 − *p*_*zz*_), yielding *P*(*x* → *z* → *y*) = *p*_*xz*_ /(1 − *p*_*xx*_) · *p*_*zy*_ /(1 − *p*_*zz*_). Note that *P*(*x* → *z* → *y*) = *P*(*x* → *z*) · *P*(*z* → *y*).

### Determining the probability of trajectories involving two groups of Boolean states

Consider a transition *c*_*j*_ → *c*_*k*_ in the SD Markov chain. We evaluate the SD Markov chain’s ability to capture the probability of trajectories of the Boolean network that correspond to this transition. Let’s introduce the notation *G*_*j*_ = π ^−1^ (*c*_*j*_) for the set of Boolean states mapped to *c*_*j*_ and *G*_*k*_ = π ^−1^ (*c*_*k*_) for the set of Boolean states mapped to *c*_*k*_. The trajectories that correspond to the *c*_*j*_ → *c*_*k*_ transition in the SD Markov chain start from any state in *G*_*j*_, only visit states within *G*_*j*_ and *G*_*k*_, and end in the first visited state in *G*_*k*_ (i.e., the trajectories do not include transitions within *G*_*k*_). We denote the aggregated probability of these trajectories **P**(*G*_*j*_ → *G*_*k*_). Here we describe and exemplify the steps of deriving this probability.

First, we use first step analysis (described above) to determine the conditional probability of each such trajectory. These trajectories include transitions within *G*_*j*_ (including self-transitions), and end with a transition from a state in *G*_*j*_ to a state in *G*_*k*_. The probabilities of the transitions within *G* are included in a |*G*_*j*_ |x |*G*_*j*_ |submatrix of the Boolean network’s transition matrix. We denote this submatrix 𝕋_1_ (*G*_*j*_ → *G*_*j*_). Analogously, the transitions that start in a state of *G* _*j*_ and end in a state of *G* _*k*_ are included in the submatrix 𝕋_1_ (*G*_*j*_ → *G*_*k*_) of dimension |*G*_*j*_ |x |*G*_*k*_ |. Thus, the system of linear equations arising from first step analysis can be expressed in a matrix form. The matrix 𝕋_∞_ (*G*_*j*_ → *G*_*k*_) of the conditional probabilities of eventually converging from each Boolean state in *G*_*j*_ to each Boolean state in *G*_*k*_ is the solution of the equation

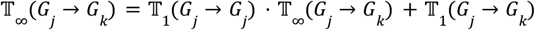

Next, **P**(*G*_*j*_ → *G*_*k*_) can be determined as a weighted sum of the elements in 𝕋_∞_ (*G*_*j*_ → *G*_*k*_) over the initial states,

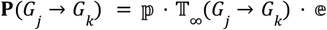

Here ℙ is a column vector of size |*G*_*j*_| that contains the initial probabilities of the states in *G*_*j*_ and 𝔼 is a row vector of size |*G*_*k*_| for which all elements are 1.

We illustrate the application of this method to determine **P**(*G*1 → *G*2) in Example **1**. The relevant submatrices of the Boolean system’s transition matrix (shown in Table 1) are

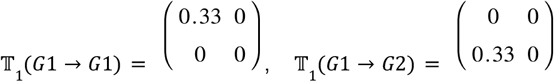

The equation for 𝕋_∞_ (*G*1 → *G*2) is

𝕋_∞_ (*G*1 → *G*2) = 𝕋_1_ (*G*1 → *G*1) · 𝕋_∞_ (*G*1 → *G*2) + 𝕋_1_ (*G*1 → *G*2). After rearranging, it becomes

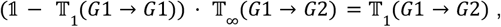

where 𝟙 is the identity matrix of dimension |*G*1| x |*G*1|. After verifying that the determinant of the matrix 𝟙 − 𝕋_1_ (*G*1 → *G*1) is not 0, we invert it and obtain

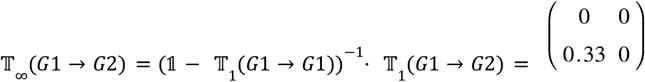

Assuming an equal initial probability distribution over the two states in *G*1,

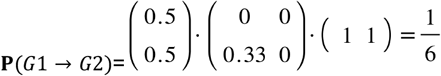

This probability is the same as the corresponding transition probability in the SD Markov chain (see Table 1).

### Determining the conditional probability of trajectories involving sequences of groups of Boolean states

A general setting for the evaluation of the SD Markov chain’s ability to capture the probabilities of sequences of events compares the probability of a path *c*_1_ → *c*_2_ → … → *c*_*n*_ in the Markov chain to the aggregated probability of trajectories that start from the set of Boolean states *G*_1_ = π^−1^(*c*_1_), go through states in *G*_1_, *G*_2_,.. in a sequence, and arrive to a state in *G*_*n*_ = π^−1^ (*c* _*n*_). Due to the sequential nature of this path,

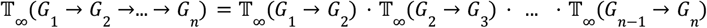

In Example 1, the conditional probability of trajectories that follow the sequence *G*1 → *G*2 → 000 satisfies

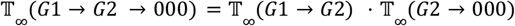

According to first step analysis,

𝕋_∞_ (*G*2 → 000) = 𝕋_1_ (*G*2 → *G*2) · 𝕋_∞_ (*G*2 → 000) + 𝕋_1_ (*G*2 → 000), which yields 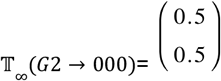

Thus, 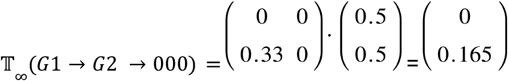

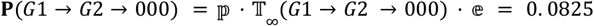

The corresponding probability in the Markov chain, *P*(*c*_1_ → *c*_2_ → 000) can be calculated in a similar way.

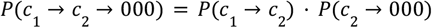

According to first step analysis, *P*(*c*_1_ → *c*_2_) = 1/6 *P*(*c*_1_ → *c*_2_) + 1/6, which yields *P*(*c*_1_ → *c*_2_) = 1/5, *P*(*c*_2_ → 000) = 1/3 *P*(*c*_2_ → 000) + 1/3, which yields *P*(*c*_2_ → 000) = 1/2.

Thus, *P*(*c*_1_ → *c*_2_ → 000) = 0. 1.

## Data availability

The datasets generated and analyzed during the current study are available in the github repository https://github.com/kyuhyongpark/sdmarkov.

## Code availability

The code for this study is available in the github repository https://github.com/kyuhyongpark/sdmarkov.

## Acknowledgements

We are grateful for the helpful suggestions of Dr. Jordan Rozum.

This study was supported by internal funds to RA from the Pennsylvania State University.

## Author contributions

KHP conceived the project, developed and implemented the algorithms, performed the comparative analyses, and wrote the manuscript. RA advised the project and wrote the manuscript. Both authors read and approved the final manuscript.

