## Supplemental Figures for "Succession-diagram-based Markov chains reveal the attractor landscape of asynchronous Boolean networks"

### Supplementary Figures

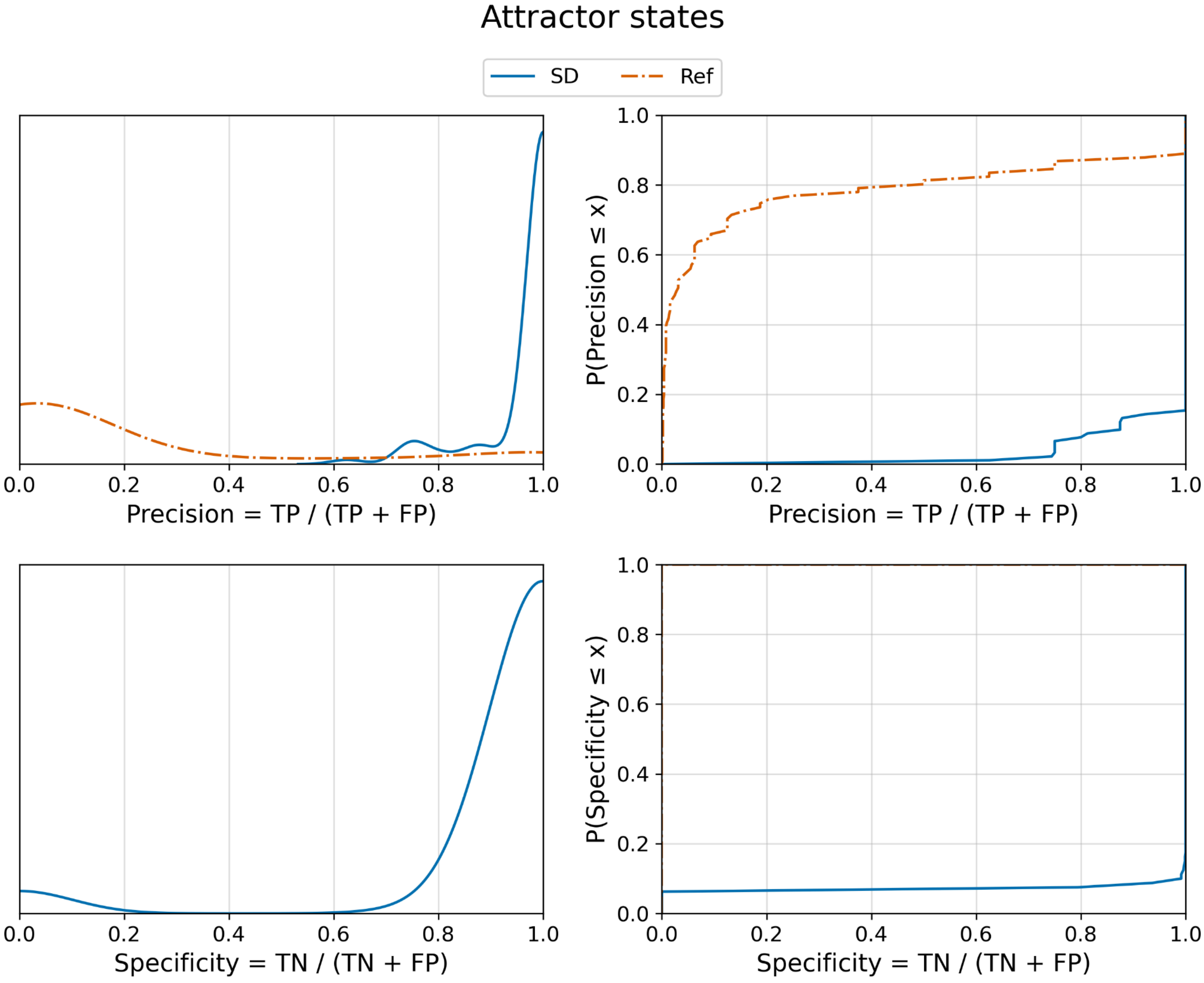


Figure **S1**: Distributions of the precision and specificity of identifying attractor states. The left panel shows the kernel density estimate (KDE) of the distributions, for which the area under each curve is equal. The right panel shows the cumulative distribution function (CDF). The precision is calculated for all 91 members of the RBN ensemble, and the specificity is calculated for the 80 RBNs that had any transient states. The reference considers that all states belong to attractors, thus its specificity is 0. The recall and NPV of the SD Markov chain is 1 for all RBNs, due to the one-to-one correspondence of attractors and minimal trap spaces. The KDE distributions show a dominant mode at 1 for the SD Markov chain, while the precision of the reference has a mode near 0.

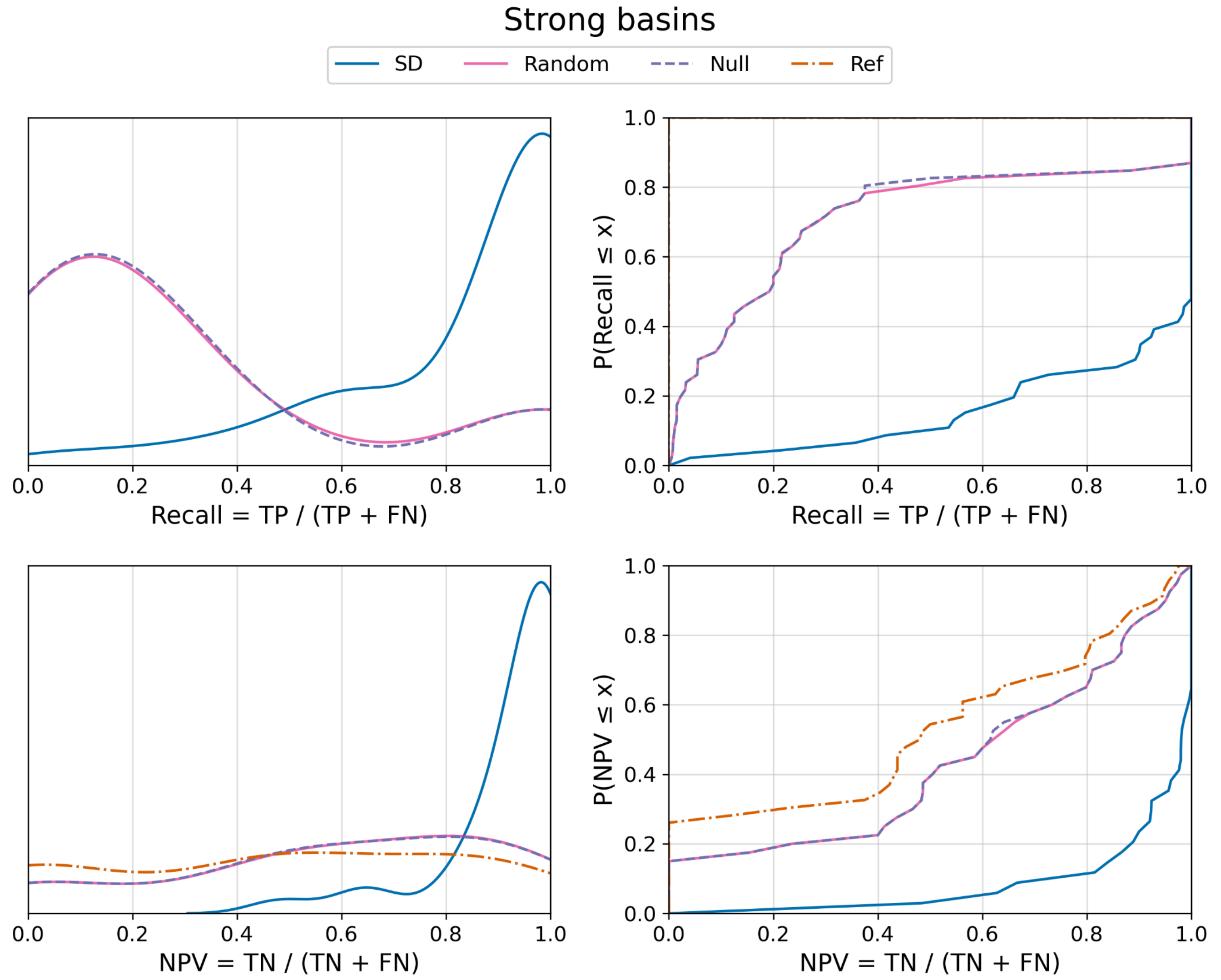


Figure **S2**: Kernel density estimate and cumulative distributions (KDE and CDF) of the recall and negative predictive value (NPV) of identifying strong basins. We use the 46 RBNs that have more than one attractor, to allow for the existence of weak basins. The reference assumes that all states belong to weak basins, thus its recall is 1 and it serves as a lower limit of the NPV. The almost perfect overlap of the continuous pink lines and dashed purple lines indicates that the Random and Null Markov chains have very close distributions. The NPV distributions of the Random and Null Markov chains are shifted toward slightly higher NPV values compared to the reference; the CDF indicates this shift by the overlapping pink and purple lines being below the brown dash-dotted line. The SD Markov chain's recall and NPV distributions are strongly skewed toward high values. In the kernel density estimates (left panels), the SD Markov chain shows a dominant mode near 1 for both recall and NPV, whereas the Random and Null Markov chains concentrate their mass at substantially lower values. This difference is further emphasized by the cumulative distributions (right panels): the SD Markov chain's cumulative distribution lies entirely below those of the Random and Null Markov chains.

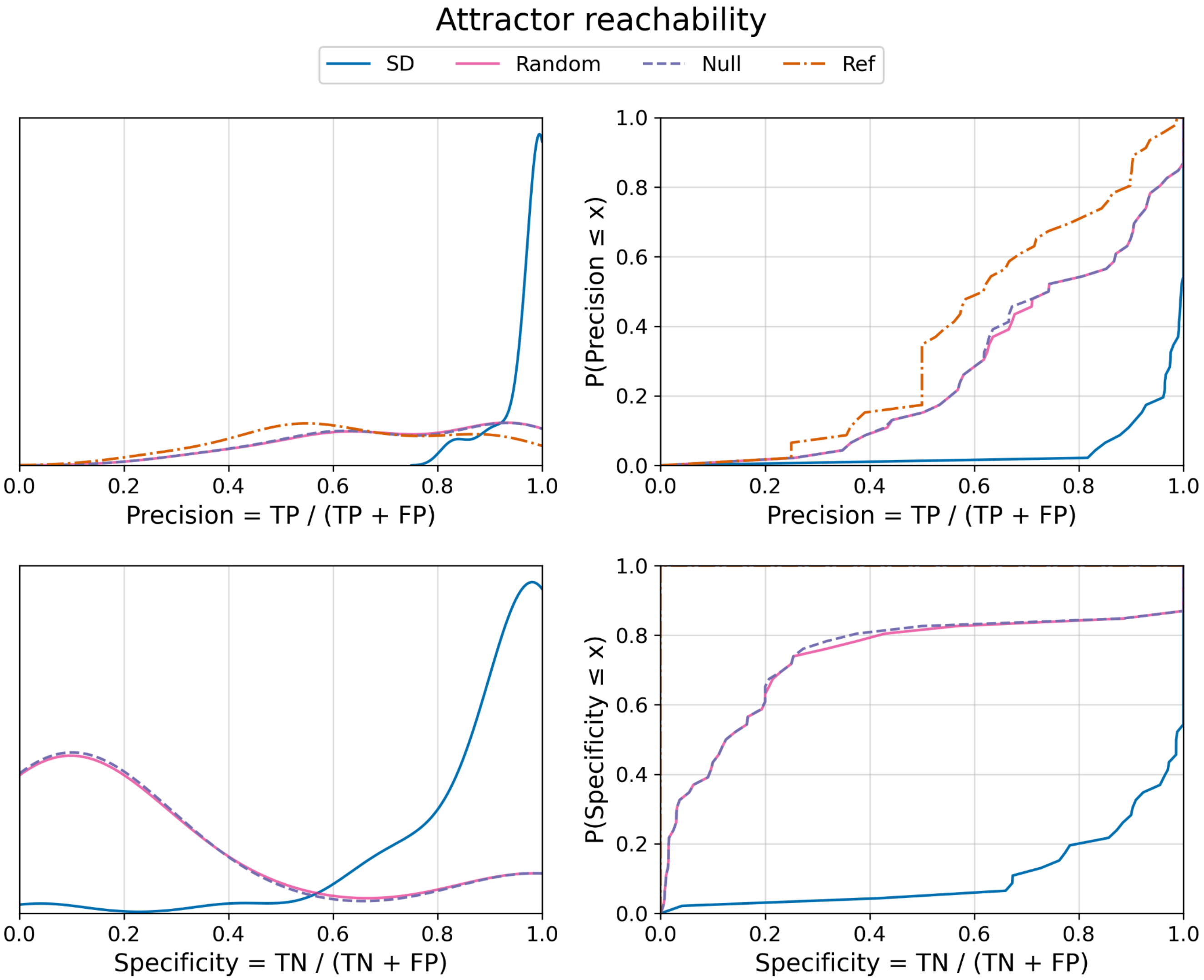


Figure **S3**: Distributions (KDE and CDF) of the precision and specificity of identifying reachable attractors from each Boolean state. We use the 46 RBNs that have more than one attractor. The reference assumes that all states reach all attractors, thus its specificity is 0 and its precision serves as a lower limit. In the KDE plots, the SD Markov chain's precision and specificity distributions have a dominant mode near 1, whereas the Random and Null Markov chains concentrate their mass at substantially lower values. The cumulative distributions of the SD Markov chain's precision and specificity lie substantially below those of the Random and Null Markov chains, and close to the theoretical best.

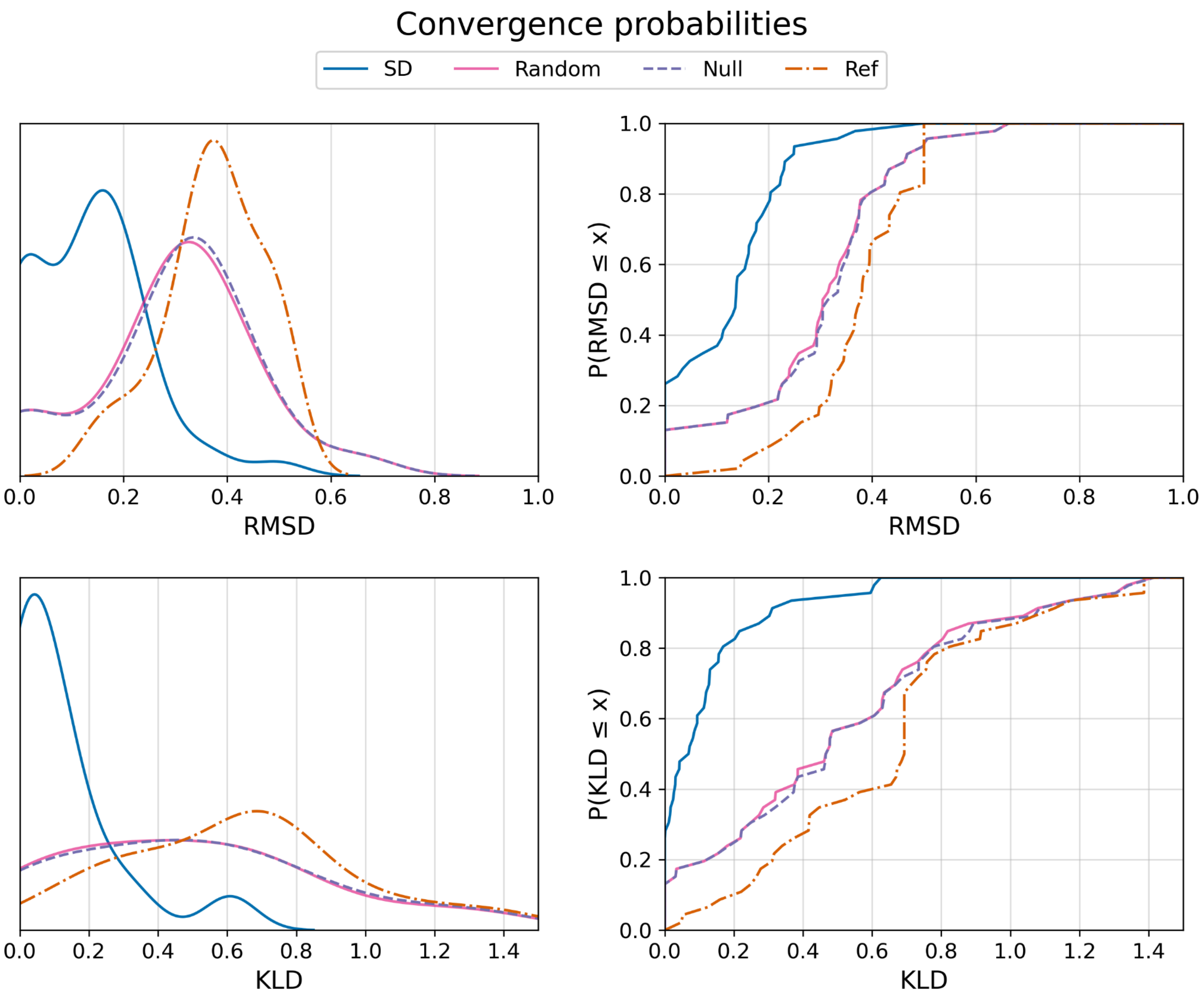


Figure **S4**: Distributions (KDE and CDF) of the RMSD and KLD between the convergence probabilities of the Boolean network and the coarse-grained Markov chains. The ideal value of RMSD and KLD is 0. We use the 46 RBNs that have more than one attractor. The reference assumes convergence to all attractors with equal probability. The SD Markov chain's RMSD and KLD values are concentrated at low values, while the mass of the Random and Null Markov chains and the reference are at intermediate values. This difference is further emphasized by the cumulative distributions: the SD Markov chain's cumulative distribution lies significantly above those of the Random and Null Markov chains and of the reference.

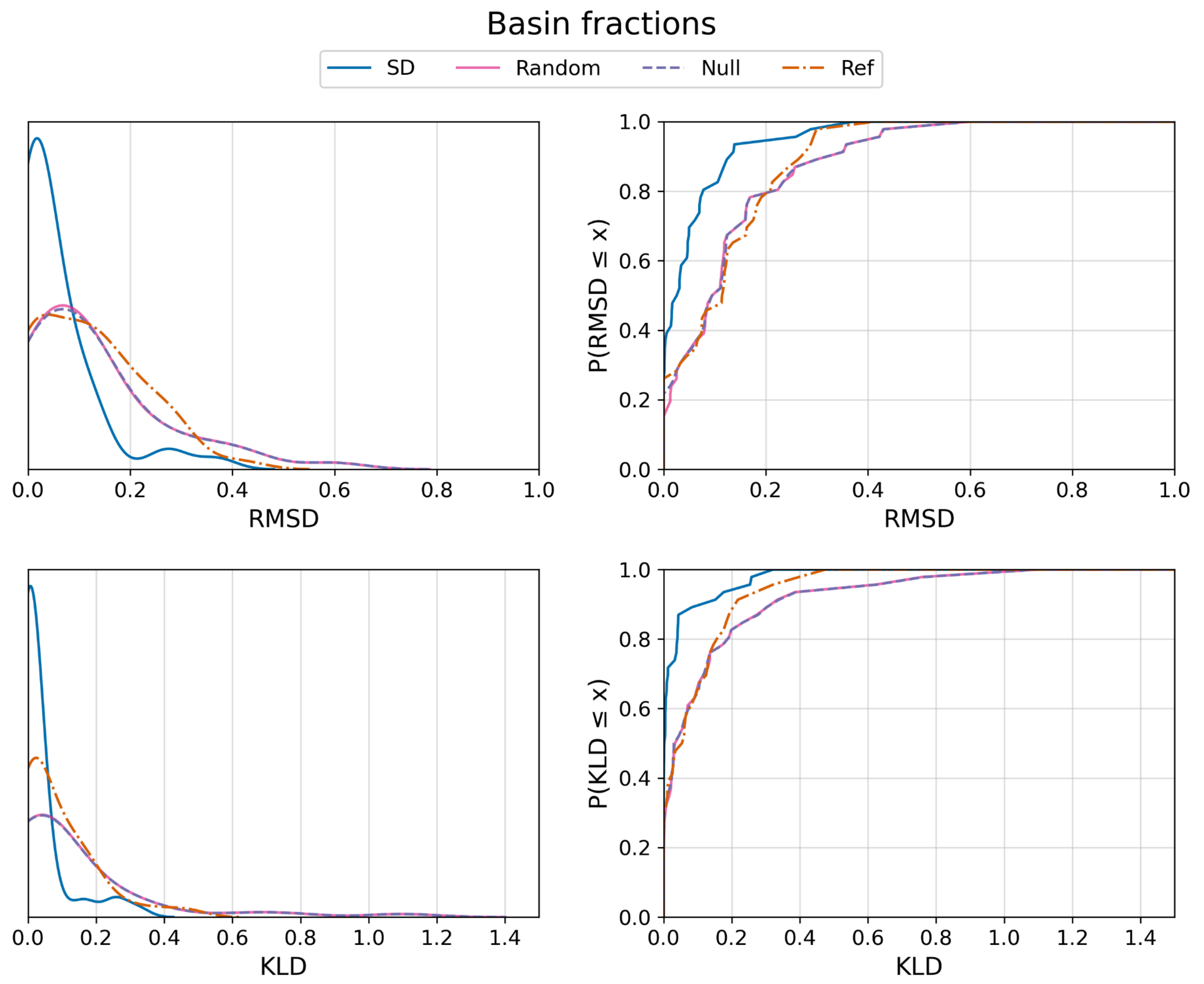


Figure **S5**: Distributions (KDE and CDF) of the RMSD and KLD between the basin fractions of the attractors of the Boolean network and the corresponding absorbing states of the coarse-grained Markov chains. We use the 46 RBNs that have more than one attractor. The reference assumes convergence to all attractors with equal probability. For both the RMSD and the KLD, the mass of the distribution of the SD Markov chain is below those of the Random Markov chains, Null Markov chain, and the reference. The cumulative distributions of the SD Markov chain lie entirely above those of the Random and Null Markov chains and of the reference.

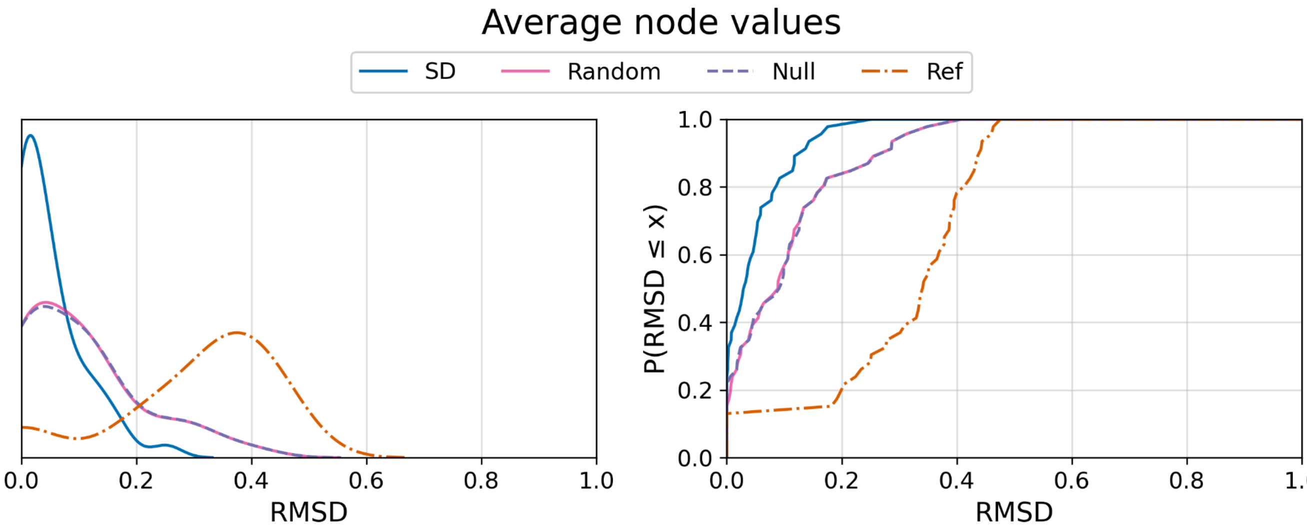


Figure **S6**: Distributions (KDE and CDF) of the RMSD between the average node values of the Boolean network and the coarse-grained Markov chains. We used the 46 RBNs that had more than one attractor. The reference assumes a value of 0.5 for every node. The mass of the RMSD distribution of the SD Markov chain is below those of the Random and Null Markov chains, and well below that of the reference. The cumulative distribution of the SD Markov chain lies significantly above those of the Random and Null Markov chains and of the reference.

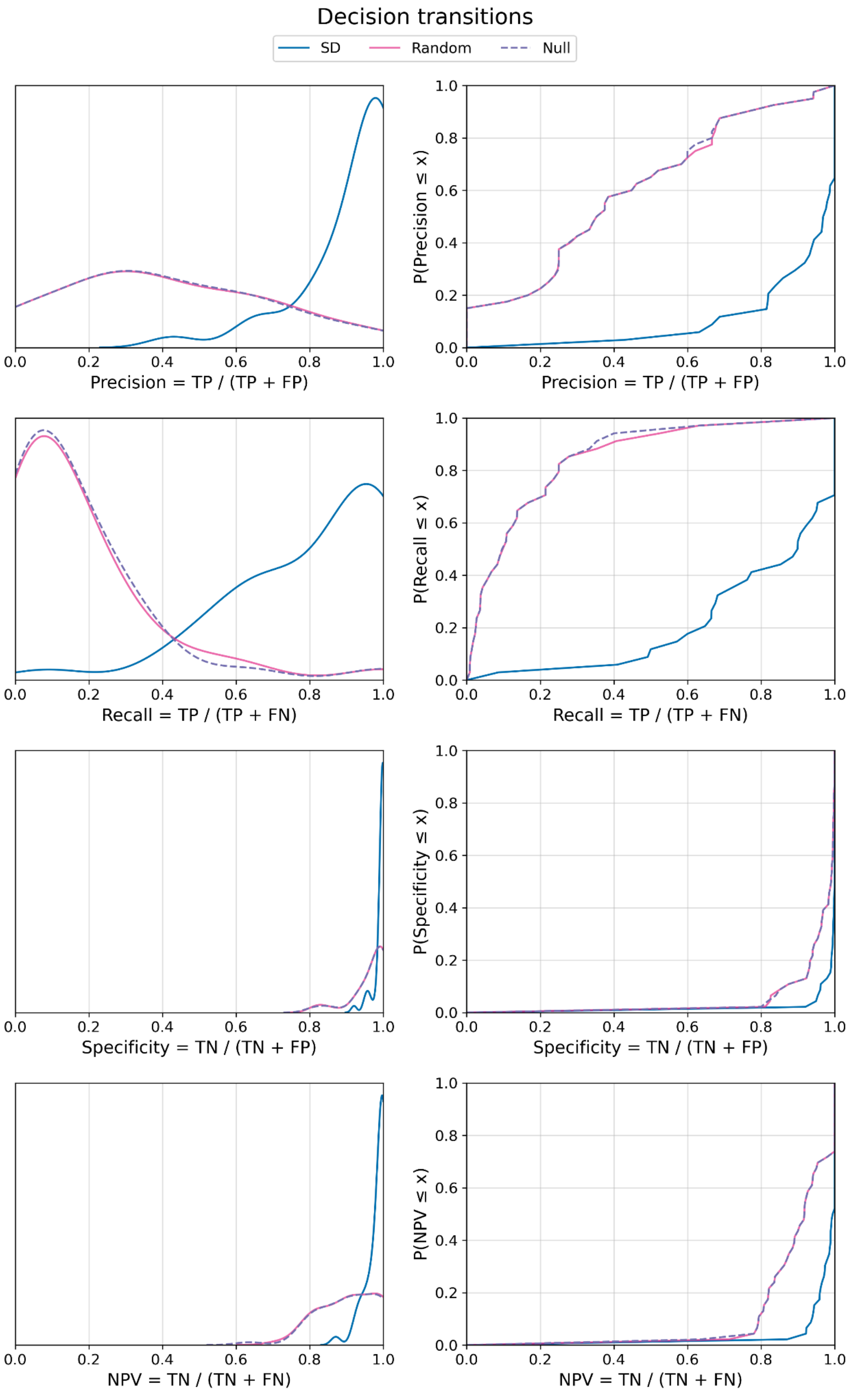


Figure **S7**: Distributions (KDE and CDF) of the precision, recall, specificity and NPV of identifying decision transitions of the Boolean network using the coarse-grained Markov chains. We use the 46 RBNs that have more than one attractor. Twelve RBNs do not have any decision transitions, which the SD Markov chain correctly identifies. The Random and Null Markov chains fail to identify 6 of these RBNs and have false positives, leading to a precision of 0 for these RBNs. The SD Markov chain's values are concentrated near 1 for all four metrics, while the Random and Null Markov chains' mass is at lower values (dramatically lower for recall). The cumulative distribution of the SD Markov chain is below those of the Random and Null Markov chains (significantly below for precision and recall) for the entire RBN ensemble.

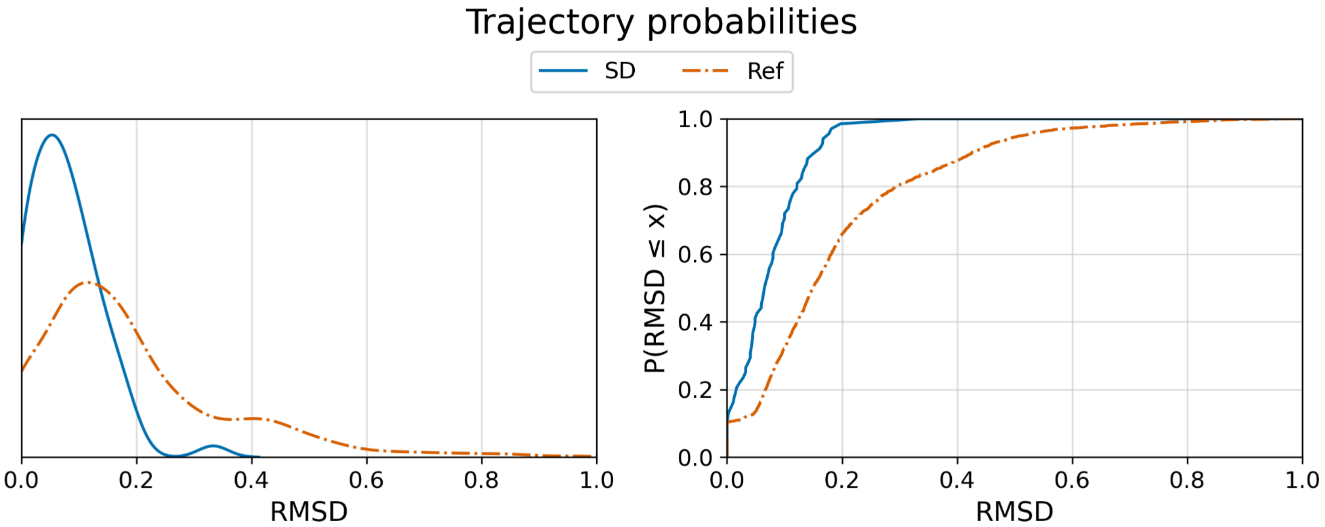


Figure **S8:** Distributions of the RMSD between the convergence probabilities over paths of length 2 or longer in the SD Markov chain and the corresponding convergence probabilities over trajectories in the Boolean network. We used the 68 RBNs whose SD Markov chains have paths of length 2. The reference captures the trajectories to the same extent as the SD Markov chain, but it randomizes the nonzero probabilities of the SD Markov chain. The mass of the SD Markov chain's RMSD is below that of the reference, and the cumulative distribution of the SD Markov chain is above that of the reference over the RBN ensemble.
